# The epigenetic regulator TRIM24 controls melanoma cell dedifferentiation and resistance to treatment in melanoma

**DOI:** 10.1101/2025.05.20.654822

**Authors:** Simon Durand, Félix Boivin, Roxane M. Pommier, Laetitia Barbollat-Boutrand, Maxime Grimont, Félix Pham, Marion Dufeu, Eric Cumunel, Raphaël Schneider, Anthony Ferrari, Anaïs Eberhardt, Stéphane Dalle, Julie Caramel

## Abstract

Cancer cell plasticity plays a key role in tumor progression and treatment resistance in melanoma. While the transcriptional programs enabling adaptative switching between melanocytic and mesenchymal phenotypes are well characterized, unravelling druggable epigenetic regulators that sustain melanoma cell adaptation and resistance remains crucial. Herein, we identified TRIM24, a bromodomain protein frequently upregulated during melanoma metastatic progression, as a crucial regulator of melanoma cell plasticity towards invasive/resistant states. shRNA-mediated knock-down of TRIM24 or degradation using a TRIM24-specific PROTAC decrease the migratory capacities and increase the sensitivity to BRAF inhibitors of melanoma cells. Integration of transcriptomic (RNA-seq) and epigenomic (ATAC-seq, CUT&Tag) analyses reveals that TRIM24 reprograms the epigenome of melanoma cells, promoting mesenchymal and repressing melanocytic transcriptional programs. We further define a TRIM24-specific transcriptional signature, that is consistently enriched in treatment-resistant mesenchymal subpopulations in melanoma single-cell RNA-seq datasets. Accordingly, analysis of TRIM24 protein expression in melanoma patients highlights that high TRIM24 expression correlates with relapse to adjuvant immunotherapy. Finally, TRIM24 knock-down in immunocompetent mouse models synergises with immune checkpoint inhibitors. Overall, our findings spotlight TRIM24 as a major epigenetic regulator driving melanoma cell dedifferentiation and resistance to therapy, representing a promising druggable target to reverse phenotype switching and resensitize to treatment.

## Introduction

Cutaneous melanoma is the most aggressive type of skin cancer. Despite drastic therapeutic advances provided by targeted therapy inhibiting BRAF/MEK and immunotherapy targeting inhibitory immune checkpoints (anti-PD-1, anti-CTLA4), 50% of cutaneous melanoma patients experience disease relapse.

Tumor cell plasticity allows rapid adaptation of tumor cells in response to exogenous stress and anti-cancer therapies, and is a major source of intra-tumor heterogeneity dramatically impacting patients’ trajectories during tumor progression and response to treatment (Marine et al., 2020). Melanoma cell plasticity involves reversible phenotypic transitions between differentiated states and undifferentiated or Neural Crest Stem Cell-like (NCSC) states with invasive properties (Hoek and Goding, 2010). By sustaining tumor cell plasticity, non-genetic reprogramming driven by transcriptional, epigenetic, or translational mechanisms supports the emergence/selection of minor but multi-resistant tumor cell subsets representing the Minimal Residual Disease responsible for disease relapse upon BRAF/MEK targeted therapy (TT) (Richard et al. 2016; Rambow et al. 2018) as well as immunotherapy (IT) (Plaschka, JITC, 2022; Pozniak et al., 2024). In particular, mesenchymal melanoma cells support immune evasion and resistance to anti-PD1, notably by preventing CD8+ T cells infiltration (Plaschka et al., 2022).

The transcriptional programs driving phenotype switching in melanoma are well characterized. Major transcriptional regulators of the melanocytic program include SOX10 and MIcrophthalmia-associated Transcription Factor (MITF), while the invasive phenotype is largely driven by AP1 complexes (Verfaillie et al., 2015). Increasing evidence indicate that the rapid transcriptional adaptations sustaining phenotype switch in response to treatment are in part epigenetically controlled, for example through EZH2-mediated rapid and reversible chromatin remodelling (Zingg et al., 2015, 2017). Interestingly, epigenetic regulators controlling melanoma cell plasticity were also associated with modulations of anti-tumor immunity (Griffin et al. 2021; Zhang et al. 2021), suggesting that epigenetic targeting could also help boost anti-tumor immune response to overcome resistance to current immunotherapies. Therefore, a better understanding of the epigenetic mechanisms underlying phenotypic plasticity and therapy resistance in melanoma may enable to develop innovative therapeutic strategies addressing treatment resistance.

Epigenetic regulators are frequently mutated or deregulated in melanoma (Halaburkova et al., 2020). In the present study, we identified *TRIM24* (*Tripartite Motif Containing 24*) /*TIF1α* as a transcriptional regulator frequently deregulated in melanoma. TRIM24 harbors a PHD-bromodomain with high affinity for histone H3, unmodified lysine 4 (H3K4me0) and acetylated lysine 23 (H3K23ac), conferring it a histone reader activity (Tsai et al., 2010). TRIM24 also possesses a RING/B-Box/Coil-coiled (RBCC) protein-protein interaction domain enabling the recruitment of co-activators and/or co-repressors to modulate gene expression (Appikonda et al. 2016).

Herein, we decipher the transcriptional and epigenetic reprogramming mediated by TRIM24 in melanoma models using CUT&Tag analyses combined with RNA-seq and ATAC-seq. Our integrative analyses define TRIM24 as a major epigenetic regulator of transcriptional programs associated with phenotypic transitions towards mesenchymal therapy-resistant state in melanoma.

## Results

### TRIM24 sustains invasive and therapy-resistant properties of melanoma cell

By exploiting TCGA analyses of epigenetic regulators alterations (Halaburkova et al., 2020), we found *TRIM24* as the most frequently altered and amplified epigenetic regulator in melanoma, before well-known regulators such as *EZH2*, *KMT2C* or *JARID2* (Fig. 1A-B). *TRIM24* is also found among top 12 genes amplified in TT or IT-resistant melanoma samples (Liu et al., 2023), although its precise role was not addressed. Moreover, *TRIM24* mRNA expression is significantly elevated in metastatic samples compared to primary tumors in the TCGA cutaneous melanoma cohort (Fig. 1C).

**Figure 1:**
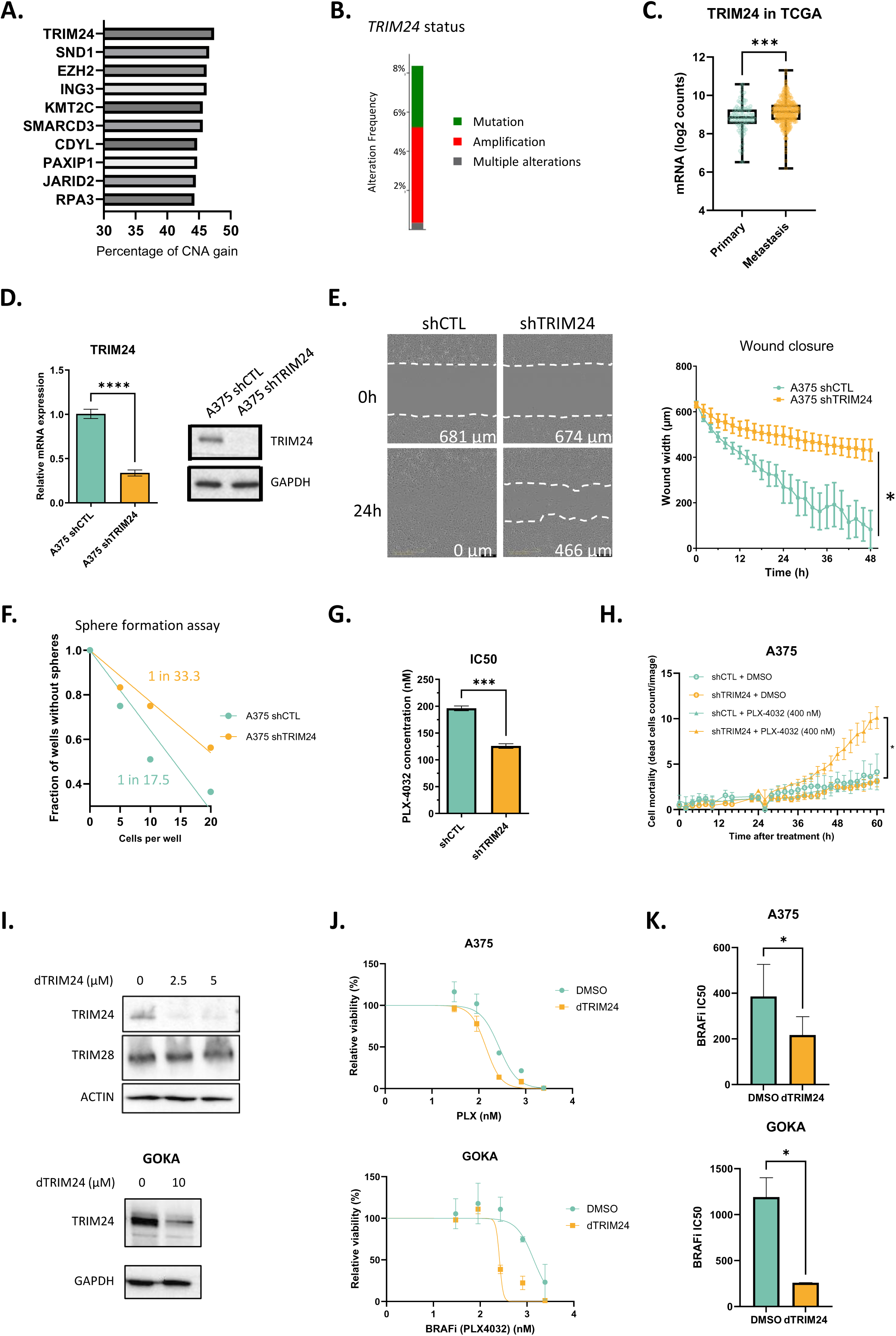
TRIM24 is deregulated in melanoma and is associated with invasive, stem-like and treatment-resistant properties of melanoma cells. **A.** Percentage of CNA gain for the 10 most frequently gained epigenetic regulators genes in the TCGA melanoma dataset. Data were extracted from Halaburkova *et al*. 2020 **B.** Frequency of genomic alterations of *TRIM24* gene in the TCGA melanoma dataset (Firehorse legacy, n = 480). **C.** *TRIM24* mRNA expression in primary and metastatic melanoma tumors from the TCGA (Firehorse legacy, n = 480). **D.** Evaluation of shRNA knockdown of *TRIM24* by RT-qPCR (left) and Western-blot (right) in A375 melanoma cells. **E.** Wound healing assay in *TRIM24* knockdown A375 cells. Representative images (left) at 0 and 24h and quantification of the wound width according to time (right) for A375-shCTL and A375-sh*TRIM24* cells. **F.** Sphere formation capacity of A375-shCTL and –sh*TRIM24.* The number of wells without spheroid at the end of the experiment for a specific starting number of cells is indicated. The fraction of cells able to form a sphere was calculated using the ELDA software (https://bioinf.wehi.edu.au/software/elda/). **G.** IC50 values for BRAFi PLX4032 in A375 after 6 days of treatment in A375-shCTL and –sh*TRIM24*. **H.** Measure of A375 cells mortality over-time using incucyte assay comparing sh*TRIM24* and shCTL cells with and without PLX4032 treatment at 400nM. **I.** Evaluation of PROTAC-mediated TRIM24 degradation using dTRIM24 assessed by Western-blot in A375 and GOKA cell lines. **J-K.** Viability curve **(J)** and IC50 values **(K)** for BRAFi PLX4032 alone or combined with dTRIM24 in A375 and GOKA cells after 6 days of treatment. P values were determined by a unpaired two-sided Mann Whitney test (**C**), a unpaired two-sided t-test (**D-E**, **G-H, K**) or by a one-way ANOVA with Tukey’s multiple comparison post-hoc test at final timepoint (**H**). Differences were considered statistically significant at *P ≤ 0.05, **P < 0.01 and ***P < 0.001. ns (non-significant) means P > 0.05.

These data prompted us to investigate the role of TRIM24 in regulating melanoma biology. For this, we generated shRNA-mediated TRIM24 knocked-down cells in two human *BRAFV600*-mutated melanoma models: A375 and C-09.10, respectively harbouring a neural-crest-like and a transitory phenotype (Fig. 1D & Supp. Fig. 1A). TRIM24 knock-down did not affect proliferation of the cells *in vitro,* nor their tumorigenic growth *in vivo* upon xenograft in nude mice (Supp. Fig. 1B-D) but was associated with major modifications of melanoma cell properties. Indeed, TRIM24 knock-down led to decreased migratory capacities of A375 melanoma cells, as evidenced by wound healing assays (Fig. 1E). Moreover, the sphere forming capacity of TRIM24-knocked-down cells in low-adherent conditions was drastically reduced, suggesting additional defects in stem-cell properties (Fig. 1F & Supp. Fig. 1E). In both A375 and C-09.10 cell lines, TRIM24-knocked-down cells exhibited increased mortality when treated with BRAF inhibitor (PLX4032) as assessed by resazurin and real-time viability assays (Fig. 1G-H & Supp. Fig. 1F-G). The BRAFi IC50 values were nearly two-fold reduced in TRIM24 knocked-down cells compared to control.

We then took advantage of a proteolysis-targeting chimera (PROTAC) degrader (dTRIM24) (Gechijian et al., 2018) to achieve specific degradation of the TRIM24 protein. We validated its specificity to degrade TRIM24 in A375 and GOKA cell lines, while not affecting TRIM28, a member of the TRIM family which shares high homology with TRIM24 (Fig. 1I). Treatment of the A375 cell line with dTRIM24 recapitulated the effects of TRIM24 knock-down, resulting in enhanced sensitivity to BRAFi treatment. Similar results were obtained in the patient-derived BRAFi-resistant melanoma cell line GOKA, where treatment with dTRIM24 drastically increased the sensitivity to BRAFi (Fig. 1J–K).

Overall, these results establish that TRIM24 loss or its pharmacological degradation in several models reverses invasive, stem-like and treatment resistance properties of melanoma cells.

### TRIM24 sustains mesenchymal melanoma cell state through major transcriptomic and epigenomic remodelling

To get deeper insights into the molecular mechanisms by which TRIM24 regulates melanoma cell properties, we first performed RNA-sequencing (RNAseq) upon *TRIM24*-knock-down. Loss of TRIM24 in the neural-crest-like/invasive A375 cell line led to major transcriptomic alterations, characterized by the repression of invasive/mesenchymal and undifferentiated signatures and the induction of melanocytic signatures (Fig. 2A-B). We then assessed whether a similar transcriptional reprogramming occurred when starting form an intermediate/transitory phenotype such as the C-09.10 cell line (Durand et al. 2024), known to harbor plastic features (Pérez-González et al. 2023). Importantly, in line with observations made in A375 cells, TRIM24 knock-down in the C-09.10 melanoma cell line induced a transcriptomic reprogramming characterized by the downregulation of invasive gene signatures and the concomitant upregulation of melanocytic differentiation programs (Supplementary Fig. 2A-B), thereby reinforcing the role of TRIM24 in promoting phenotype switching across both invasive and transitory melanoma states. This reprogramming was further substantiated by the modulation of canonical lineage markers, as TRIM24 depletion led to increased expression of MITF and SOX10, hallmarks of the melanocytic state, along with a reduction in mesenchymal and undifferentiated markers, including ZEB1 and SOX9 (Supplementary Fig. 2C–D).

**FIGURE 2:**
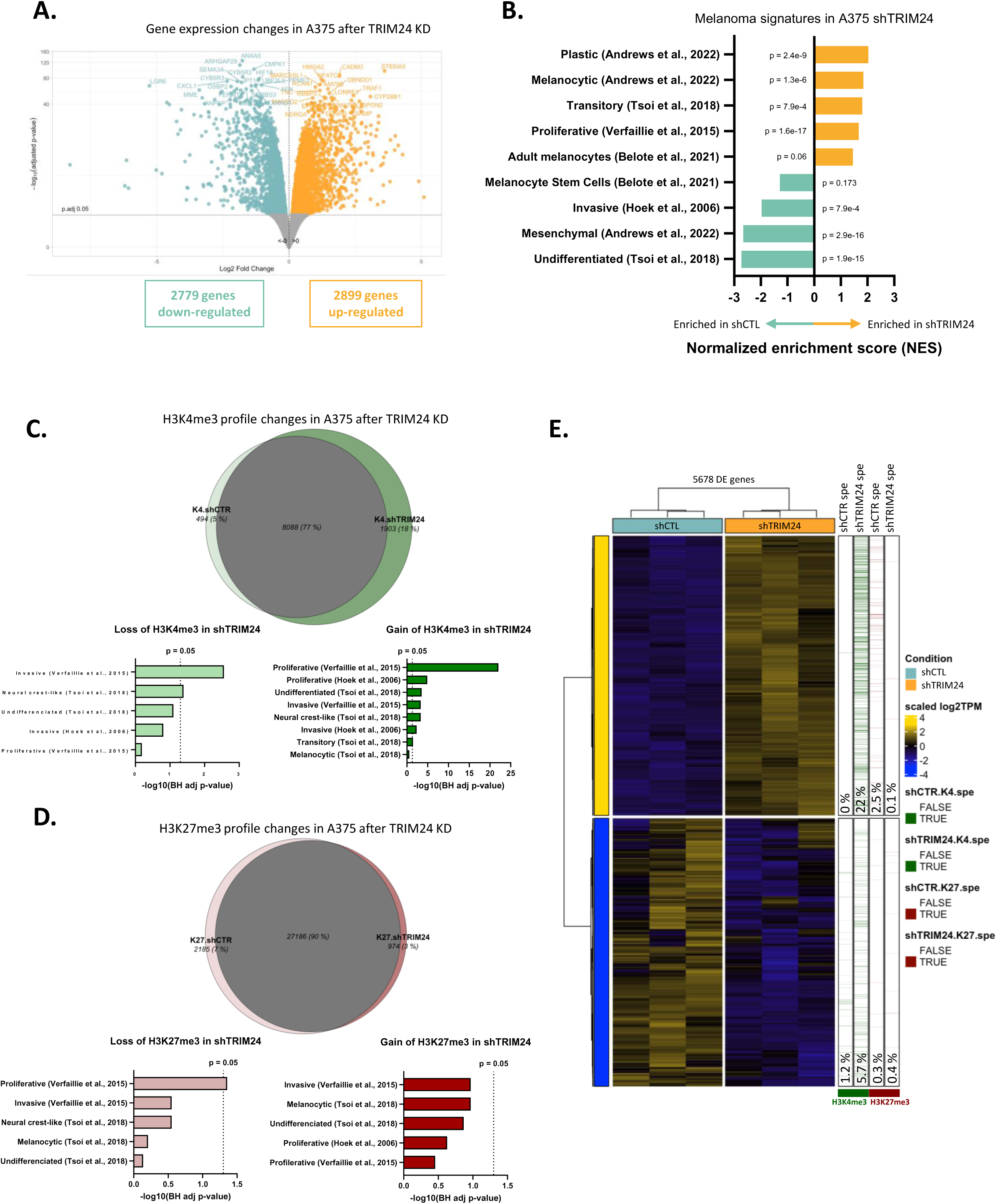
*TRIM24* concomitantly promotes invasive and represses proliferative transcriptional programs through major epigenetic remodelling. **A.** Volcano plot of differentially expressed (DE) genes identified by RNA-seq in A375 after *TRIM24* knock-down (KD). **B.** Normalized enriched scores (NES) upon *TRIM24-*KD for melanoma cell phenotype signatures. **C-D.** Venn diagrams of CUT&Tag data displaying the differential binding of the activating H3K4me3 (**C**) or the repressive H3K27me3 (**D**) histone mark associated to a gene promoter in A375-sh*TRIM24* versus –shCTL cells. Melanoma cell signatures enriched in genes presenting a reduction or a gain of these histone marks are scored. **E.** Heatmap of DE genes upon *TRIM24-*KD in A375 cells. H3K4me3 and H3K27me3 histone marks specific to sh-*TRIM24* or shCTL are indicated. The percentages of genes presenting a signal for the indicated histone marks, among the sh*TRIM24*-activated (top) or sh*TRIM24*-repressed (bottom), are indicated.

Consistent with these findings, analysis of the TCGA melanoma cohort revealed a positive correlation between *TRIM24* expression and neural crest-associated markers *NGFR* and *ZEB1*, and a negative correlation with the differentiation markers *MITF* and *SOX10* (Supplementary Fig. 2E). Stratification of tumors based on *TRIM24* expression levels further demonstrated that TRIM24^high^ tumors were significantly enriched for neural crest-like (NCL) signatures and displayed reduced melanocytic differentiation signatures, supporting a functional association between *TRIM24* expression and maintenance of mesenchymal melanoma cell states (Supplementary Fig. 2F).

Next, to characterize the epigenome reconfiguration associated with TRIM24-mediated transcriptional reprogramming, the pattern of histone modifications was analyzed by CUT&Tag for both active (trimethyl-H3K4) and repressive (trimethyl-H3K27) histone marks in A375 and C-09.10 cells (Fig. 2C-E & Supp. Fig. 3). Interestingly, *TRIM24* knock-down induced an opposite modification of active *vs* repressive histone marks, with increased H3K4me3 and decreased H3K27me3 signals. Consistently with RNA-seq results, *TRIM24* knock-down caused a specific enrichment of H3K4me3 marks on proliferative and melanocytic gene promoters, while this permissive epigenetic modification was depleted on invasive and NCL genes (Fig. 2C & Supp. 3A-B). An opposite effect was observed with H3K27me3, the repressive mark being significantly depleted on proliferative gene promoters (Fig. 2D & Supp. Fig. 3C-G). Analyses of histone methylation patterns on differentially expressed genes (DEGs) confirmed the link between TRIM24-mediated epigenetic reprogramming and associated transcriptomic changes (Fig. 2E). Importantly, 22% of upregulated genes showed an increase in H3K4me3 upon *TRIM24* knock-down. Conversely, most of the H3K27me3 peaks present on upregulated genes were lost upon *TRIM24* knock-down.

Overall, we demonstrate that *TRIM24* expression induces a profound chromatin remodeling associated with transcriptional reprogramming from melanocytic towards mesenchymal programs.

### TRIM24 binds to promoter and enhancer regions associated with specific histone 3 K23ac pattern

To further elucidate TRIM24 mechanisms of action at the chromatin level, CUT&Tag analyses of TRIM24 binding profile were performed in the A375 and C-09.10 cell lines. The majority of TRIM24 binding peaks in A375 were located in promoter regions (51%), centered on the transcriptional start site (Fig. 3A-B). Of note, 39% of TRIM24-occupied sites were observed in introns and intergenic regions. When looking at promoters-associated peaks, most of TRIM24 peaks observed in A375 were also present in C-09.10 (n=4026) (Fig. 3C and Supp. Fig. 4A).

**FIGURE 3:**
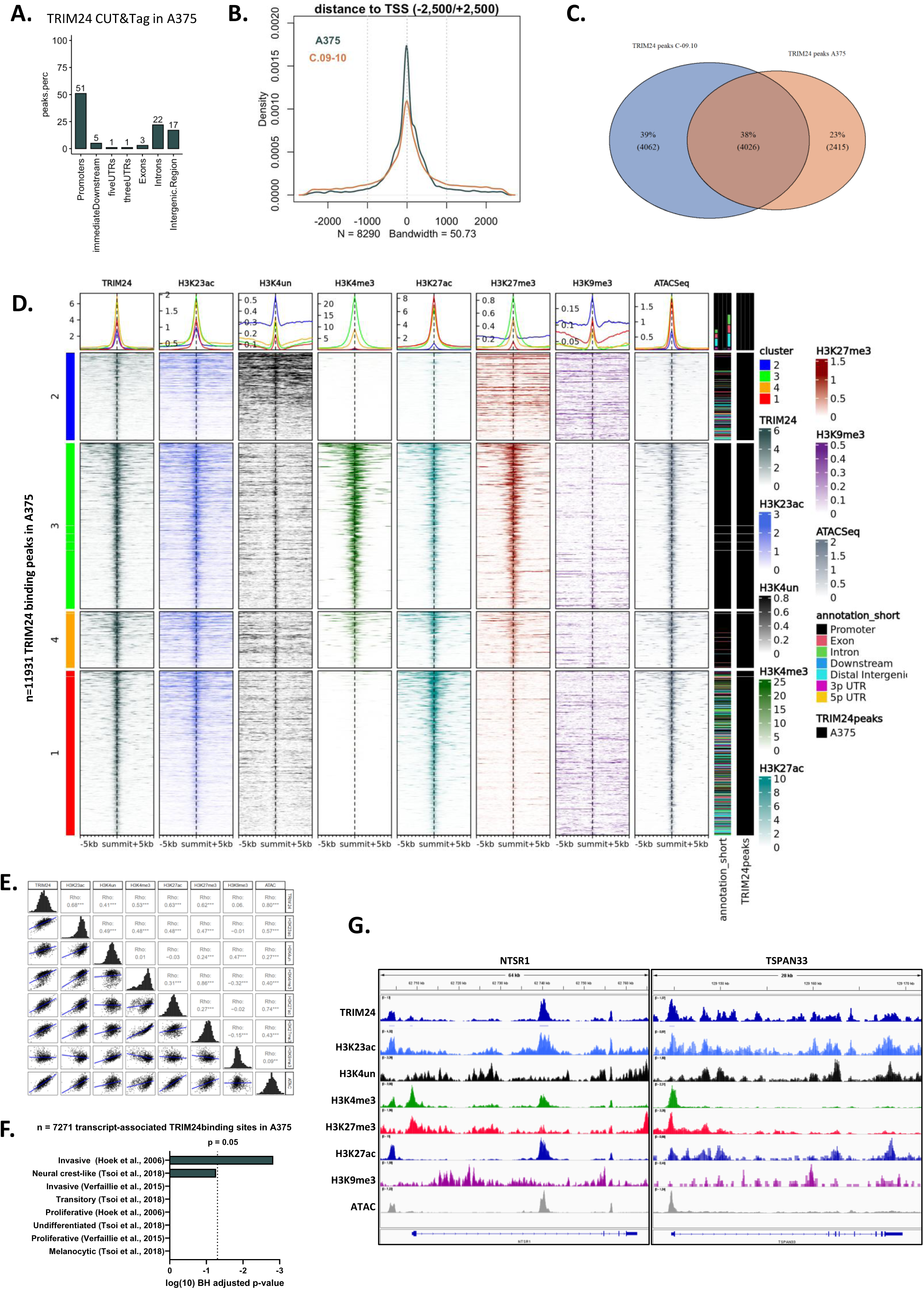
TRIM24 binds to promoter and enhancer regions of genes associated with melanoma cell state through recognition of specific histone patterns. **A.** Distribution of TRIM24 peaks across the genome of A375 cells identified by TRIM24 CUT&Tag analysis. **B.** Distance of TRIM24 peaks to the transcription start site in A375 (green) and C-09.10 (orange) cells. **C.** Venn diagram representing the overlap of TRIM24 binding peaks in A375 and C-09.10 cells. **D.** Read density heatmap of peak profiles for histone marks (H3K23ac, H3K4un, H3K4me3, H3K27ac, H3K27me3 and H3K9me3) relative to summits of TRIM24 peaks and chromatin accessibility (ATAC-seq) in A375 cells. The profiles are clustered using hierarchical clustering method, the profile intensity of each mark in each cluster is indicated. Genome annotations are indicated. **E.** Correlogram studying the association between TRIM24, histones marks and ATAC-seq peaks. **F.** Enrichment of melanoma cell phenotype signatures on genes associated to a TRIM24 binding site in A375 cells. **G.** IGV visualisation of *NTSR1* and *TSPAN33* loci, belonging respectively to the invasive and the proliferative melanoma signatures (Verfaillie et al., 2015). The binding and the peak calling of TRIM24 as well as the H3K23ac, H3K4un, H3K4me3, H3K27me3, H3K23ac, H3K9me3 and ATAC signals are included. Two replicates are overlayed for each condition.

We next performed additional CUT&Tag analyses for histone marks described in the literature as associated with TRIM24 binding, namely unmethylated H3K4 (H3K4un) and acetylated-H3K23 (H3K23ac) (Tsai et al., 2010), as well as histones marks well-known for governing gene expression such as H3K27ac (enhancers) and H3K9me3 (heterochromatin) (Talbert and Henikoff, 2021). We also analyzed chromatin accessibility by ATAC-seq to perform integrative analysis of chromatin alterations correlating with TRIM24 binding profile. TRIM24 binds to accessible regions of both gene promoters and distal enhancers, and co-locates specifically with H3K23ac histone marks as evidenced by the strongest correlation score (Fig. 3D-E and Supp. Fig. 4B-C). However, in contrast to previous studies in other cancer types (Tsai et al., 2010), TRIM24 binding did not correlate with H3K4un profile in melanoma cells. In contrast, H3K4un was rather associated with heterochromatin regions exhibiting enriched H3K9me3 signal (cluster 2). Importantly, TRIM24 binding was not only observed in gene promoters close to the TSS enriched for H3K4me3 marks (clusters 3 and 4), but also in enhancers regions, introns, and distal intergenic regions presenting an enriched H3K27ac3 signal (cluster 1) (Fig. 3D-E). Consistent with its role in sustaining invasive melanoma cell states, TRIM24 showed preferential binding to genes of invasive signature (Verfaillie et al. 2015) (Fig. 3F), as exemplified on the promoter of the *NTSR1* gene (Fig. 3G). Moreover, TRIM24 binding is also observed on genes from the proliferative signature, as shown on the *TSPAN33* gene promoter (Fig. 3G). As evidenced with these two genes, TRIM24 binding profile shares extensive similarity with H3K23ac signal but not with H3K4un.

Altogether, our findings delineate a unique chromatin-binding landscape of TRIM24 in melanoma, preferentially associating with accessible chromatin at promoter and enhancer regions marked by H3K23ac and directly locates on genes defining melanoma cell states.

### TRIM24 binds to promoter regions of melanoma cell states genes poised for transcriptional activation or repression

We next investigated the epigenetic remodeling of genes deregulated upon *TRIM24* knock-down. Notably, we observed a significant enrichment of TRIM24 binding among these genes, with 41% exhibiting TRIM24 binding at their promoters (Fig. 4A & Supp. Fig. 5A). TRIM24 binding was similarly distributed between upregulated and downregulated genes, with respectively 40% of upregulated and 42% of downregulated genes upon *TRIM24* knock-down harboring TRIM24 binding sites (Fig. 4A). Upon *TRIM24* knock-down, upregulated TRIM24-bound genes were significantly enriched for proliferative and transitory melanoma signatures, whereas downregulated TRIM24-bound genes were associated with invasive and Neural-Crest/undifferentiated transcriptional programs (Fig. 4B & Supp. Fig. 5B).

**FIGURE 4:**
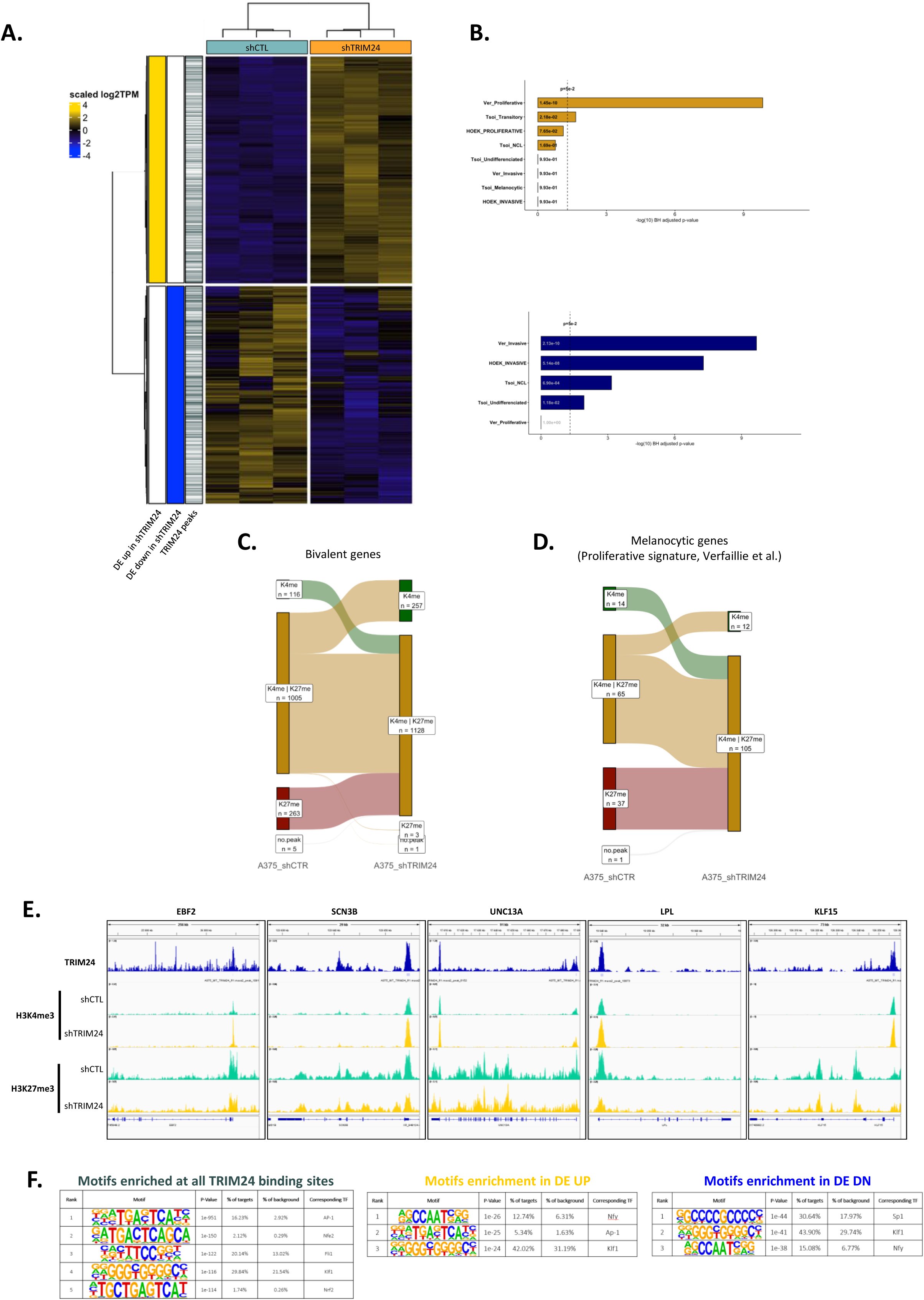
TRIM24 dualy binds to promoter regions associated with transcriptional activation or repression and alters the chromatin status of bivalent genes. **A.** Heatmap displaying the expression of differentially expressed genes upon *TRIM24-*KD in A375 cells. The binding of TRIM24 to these genes is indicated. **B.** Enrichment of melanoma cell phenotype signatures on genes associated with a TRIM24 peak and differentially expressed upon *TRIM24-*KD in A375 cells for upregulated (top) and downregulated (bottom) genes. **C.** Effect of *TRIM24* loss on genes presenting a bivalent status in A375 cells. Evolution of H3K4me3 and H3K27me3 histone marks patterns upon *TRIM24-*KD on bivalent genes. **D.** Evolution of H3K4me3/H3K27me3 marks on bivalent genes composing the melanoma proliferative signature (Verfaillie et al., 2015). **E.** IGV visualisation of *EBF2*, *SCN3B*, *UNC13A*, *LPL*, *KLF15* loci presenting a bivalent status. The binding of TRIM24 as well as H3K4me3 and H3K27me3 signals upon *TRIM24-*KD are included. Two replicates are overlayed for each condition. **F.** Top 5 HOMER-identified enriched DNA binding motifs at all TRIM24 binding sites (left), and at genes up-regulated (middle) and down-regulated (right) upon *TRIM24-*KD in A375 cell, when considering a 200 bp region. Associated p-values, percentages of motif representation on target and background signal are indicated.

We then focused on genes exhibiting a bivalent chromatin profile, marked by both active (H3K4me3) and repressive (H3K27me3) histone modifications, which are known to poise genes for rapid chromatin state transitions during differentiation (Bernstein et al. 2006). TRIM24 binding was enriched at such bivalent gene loci (Supp. Fig. 5C). Interestingly, over 40% of bivalent genes exhibited altered chromatin states upon *TRIM24* knock-down, with the majority shifting towards a more open and transcriptionally permissive configuration (Fig. 4C & Supp. Fig. 5D). Chromatin opening was confirmed among bivalent genes associated with the proliferative melanoma signature (Verfaillie et al. 2015) (Fig. 4D & Supp. Fig 5E). For example, TRIM24 binding in promoters associated with the acquisition of a more active chromatin state upon *TRIM24* knock-down was confirmed in several genes of melanoma melanocytic signature (*EBF2*, *SCN3B*, *UNC13A*, *LPL*, *KLF15*), characterized by their important role in neural system development (Fig. 4E & Supp. Fig. 5F).

As TRIM24 recognizes histone marks rather than genomic sequences, we hypothesized that its effect on transcription might differ depending on the recruitment of co-activators and/or co-repressors. To identify transcription factors (TFs) that may cooperate with TRIM24, we performed DNA-binding motifs enrichment analysis on TRIM24-occupied sites with HOMER. Motif enrichment analysis at TRIM24 binding sites evidenced an enrichment for the Activator Protein-1 (AP-1) complex members JUN and FOS, major regulators of the mesenchymal state (Verfaillie et al., 2015), in both A375 and C-09.10 cells (Fig. 4F & Supp. Fig. 5G). Other predicted TFs included members of the Specificity protein/Krüppel-like factor (Sp/KLF) family which are deregulated in cancer and melanoma (Tetreault et al., 2013; Huh et al., 2010). Of note, the AP-1 complex motif was more specifically enriched in the upregulated than in the downregulated genes (Fig. 4F), suggesting a dual and selective recruitment of co-activators and co-repressors by TRIM24 in melanoma cells.

Altogether, integrative transcriptomic and epigenomic analyses reveal a critical role for TRIM24 in orchestrating a dynamic transcriptional reprogramming through localized and selective epigenetic remodeling at target loci. TRIM24 modulates chromatin accessibility and transcriptional activity at developmentally regulated bivalent genes contributing to melanoma phenotype plasticity, by activating neural crest-like and invasive gene programs while repressing melanocytic and proliferative signatures.

### A TRIM24-associated gene signature is enriched in IT-resistant mesenchymal cells in melanoma public datasets

We then took advantage of our unique TRIM24-related RNAseq dataset to define a melanoma-specific TRIM24 signature. To do so, we selected the intersection of genes differentially expressed upon *TRIM24* knock-down in C-09.10 and A375 cells, resulting in a signature composed of 42 positively and 62 negatively TRIM24-regulated genes (Fig. 5A).

**FIGURE 5:**
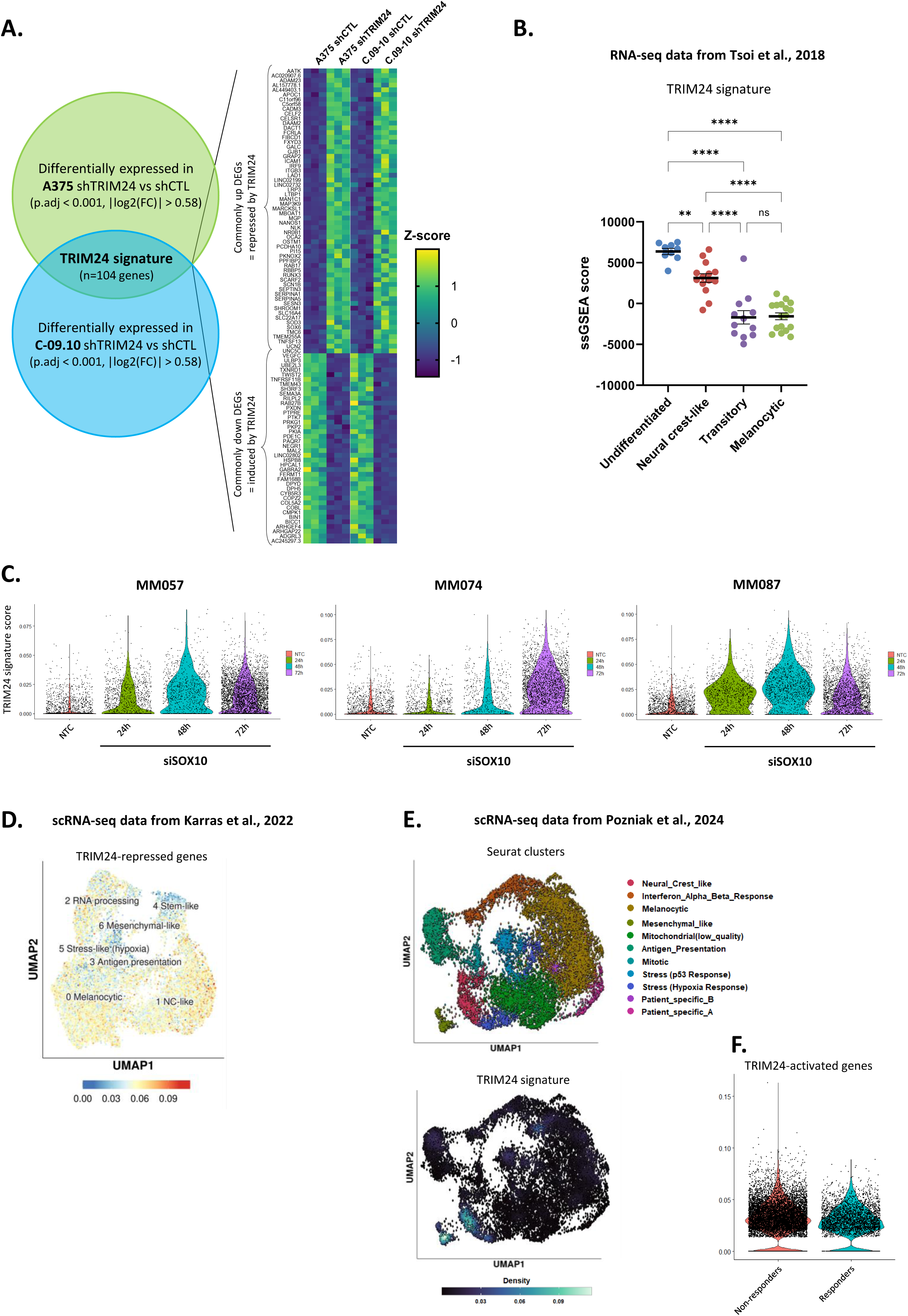
TRIM24 signature is enriched in mesenchymal and NCL therapy-resistant melanoma cell states from patients and correlates with resistance to IT. **A.** Establishment of TRIM24 signature by combining DE genes in A375 and C-09.10 upon *TRIM24*-KD. **B.** ssGSEA score of TRIM24 signature in melanoma cell lines dataset from Tsoi *et al*. **C.** Violin plot of TRIM24 signature score in single-cell RNA-seq data from 3 melanoma short term cultures transfected with SOX10 siRNA or non-targeting control (NTC) from Wouters *et al*. **D.** Featureplot displaying TRIM24 signature score in murine melanoma single-cell RNAseq data from Karras *et al*. **E.** Featureplot displaying TRIM24 signature score in human metastatic human melanoma tumor single-cell RNAseq data from Pozniak *et al*. **F.** TRIM24 signature according to the ICB response status in patients from the Pozniak *et al*. dataset. P values were determined by a one way ANOVA test with a Tukey correction (**B**). Differences were considered statistically significant at *P ≤ 0.05, **P < 0.01 and ***P < 0.001. ns (non-significant) means P > 0.05.

We used this melanoma-specific TRIM24 signature to explore public RNAseq and scRNAseq melanoma datasets to assess TRIM24 correlation with melanoma cell states and resistance to therapy. In RNA-seq data from melanoma cell lines with well-characterized phenotype (Tsoi et al., 2018), TRIM24 signature was highly increased in undifferentiated and neural-crest-like cell lines as compared to transitory and melanocytic ones (Fig. 5B). Moreover, in a larger panel of melanoma cell lines from the CCLE, the TRIM24 signature strongly correlates with the invasive signature and anti-correlates with the melanocytic one (Supp. Fig. 6A). TRIM24 signature was also progressively gained in short-term cultures upon switch towards mesenchymal state induced by *SOX10* knock-down (Fig. 5C) (Wouters et al., 2020). Consistently, the activity of the SOX10 transcription factor is enriched in the TRIM24-repressed genes (Supp. Fig. 6B).

Using this *in vitro* validated signature, we next assessed its relevance *in vivo* in scRNAseq datasets. Genes repressed by TRIM24 were highly depleted from mesenchymal and stem-like subpopulations in scRNAseq from a murine melanoma model (Karras et al., 2022) (Fig. 5D & Supp. Fig. 6C), and TRIM24 signature was selectively enriched in mesenchymal and NCL subpopulations from scRNAseq of human metastatic melanoma tumors (Pozniak et al., 2024) (Fig. 5E). Importantly, TRIM24 signature was upregulated in samples from IT-resistant patients (Fig. 5F & Supp. Fig. 6D). Genes repressed by TRIM24 were associated with improved response to IT in on-treatment biopsies (Supp. Fig. 6E).

Overall, we defined a melanoma-specific TRIM24 signature and further demonstrate its enrichment in mesenchymal melanoma sub-populations resistant to IT in patient samples.

### TRIM24 drives resistance to immunotherapy in melanoma

To further investigate TRIM24 expression pattern in melanoma patient lesions, and its association with treatment response, we performed multi-immunofluorescence staining in a cohort of stage III melanoma patients treated with adjuvant anti-PD1 immunotherapy stratified according to their relapsing status at 1 year following treatment initiation (n=52, n=17 relapsing and n=35 non-relapsing patients) (Supplementary Table 1) (Fig. 6A). Whole tumor analysis revealed higher TRIM24 protein expression in the tumor as compared to the stroma, highlighting its tumor restricted expression (Fig. 6B-C). Furthermore, intra-tumoral heterogeneity of TRIM24 expression was observed, with regions displaying drastically increased expression (Fig. 6D). By stratifying patients into TRIM24^high^ (top 25%) or TRIM24^low^ (bottom 75%) (Supp. Fig. 7), we found that an elevated TRIM24 protein level is associated with relapse under anti-PD-1 immunotherapy in this cohort. A significant enrichment in relapsing patients was found among TRIM24^high^ melanoma samples (Fig. 6E). Accordingly, TRIM24^high^ patients presented a statistically significant two-fold worse relapse-free survival following adjuvant immunotherapy (Fig. 6F).

**FIGURE 6:**
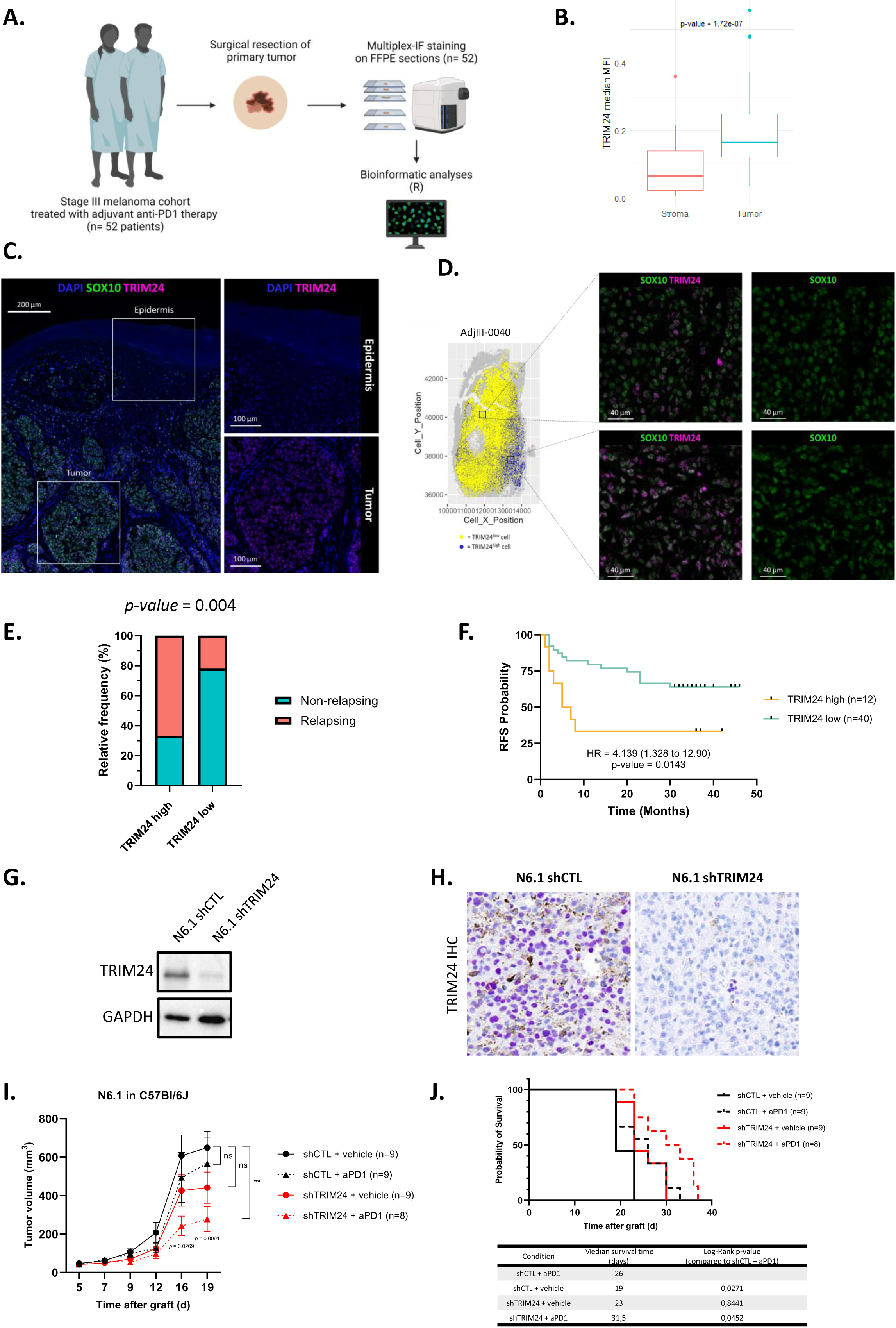
Increased TRIM24 expression is associated with resistance to immunotherapy in melanoma. **A.** Schematic multiplex-IF workflow in a cohort of stage III melanoma patients treated with adjuvant anti-PD1 therapy (n=52). **B.** Boxplot showing the immunofluorescence intensity of TRIM24 protein in the stroma and in the tumor (n = 52). The p-value was calculated by unpaired two-sided Mann-Whitney test. **C.** Representative image of a multiplex-IF on a melanoma tumor showing DAPI (blue), SOX10 (green) and TRIM24 (pink) labeling. The inserts show TRIM24 signal in the epidermis and in the tumor area. **D.** Reconstruction of a representative heterogeneous tumor as whole slide. Each dot represents one cell. The color reflects TRIM24 expression level above a threshold value: yellow (low) and blue (high). **E.** Frequency of relapsing and non-relapsing patients in the TRIM24^high^ (n=12) and the TRIM24^low^ (n=40) groups. Chi-squared test. **F.** Kaplan– Meier relapse-free survival (RFS) probability curves for melanoma patients segregated in TRIM24^high^ and TRIM24^low^ groups. HR denotes hazard ratio. **G-H.** Efficiency of shRNA-mediated *TRIM24* knock-down assessed by Western Blot (**G**) at the day of injection and by IHC (**H**), purple chromogen) at the end of the experiment. **I-J.** Tumor growth (**I**) and survival probability with log-rank test (**J**) of shCTL and sh*TRIM24* N.6.1 cells injected in C57BL6/J mice treated with anti-PD1 or vehicle. n=9 per group. P values were determined by Wilcoxon matched-pairs signed rank test (**B**), Chi-squared test (**E**) and a log-rank (Mantel–Cox) test (**F**). I. Differences were considered statistically significant at *P ≤ 0.05, **P < 0.01 and ***P < 0.001. ns (non-significant) means P > 0.05.

The association of TRIM24 expression with relapse under IT prompted us to evaluate the putative synergistic effect of TRIM24 knock-down in combination with anti-PD1. To do so, we generated TRIM24-knocked-down cells from a syngeneic mouse model, that we already described as partially responding to anti-PD1 (N6.1) (Plaschka et al., 2022) (Fig. 6G). TRIM24 knock-down was confirmed in the tumors at the protein level by IHC (Fig. 6H). While associated with a mild reduction of tumor growth in untreated mice, TRIM24 knock-down strongly potentiates response to anti-PD1 (Fig. 6I) and improves mice survival (Fig. 6J). Together, this demonstrates that TRIM24 expression is sufficient to drive resistance to anti-PD1 treatment in mice.

Together, our results support a central role for TRIM24 in preventing IT efficacy through its ability to promote mesenchymal/treatment-resistant melanoma cell states.

## Discussion

Epigenetic regulators have been proposed to account for the rapid adaptation of melanoma cells to treatment but their precise role in phenotype switching and immune escape still need to be deciphered. Our data identify TRIM24 as a novel major regulator of melanoma phenotype plasticity and resistance to therapies. We demonstrate that TRIM24 is required for maintaining mesenchymal and NCL cell states, and find that high expression of TRIM24 is associated with resistance to immunotherapy and unfavorable prognosis in melanoma patients.

Our results show that TRIM24 orchestrates the plasticity of melanoma cells, with dual activity sustaining an undifferentiated/NCL phenotype while repressing a melanocytic/proliferative program. Our integrative analyses of RNA-seq, ATAC-seq and CUT&Tag data for TRIM24 and histone marks allowed to decipher how TRIM24 transcriptional complex epigenetically reprograms melanoma cells. We show that TRIM24, which is a non-DNA binding epigenetic regulator, reads the local acetylation state of H3K23, but not H3K4un, in chromatin accessible regions. TRIM24 is physically located in gene promoters and more distal enhancers and is significantly enriched in the vicinity of genes regulating melanoma cell phenotype. TRIM24 recruitment induces an epigenetic remodelling and exerts a dual effect on gene transcription, with concomitant enhancement of a NCL/invasive gene program and repression of the melanocytic/proliferative gene program. This epigenetic and transcriptomic reprogramming finally induces a switch in melanoma cell properties. Our results also highlight a novel role for TRIM24 in regulating chromatin dynamics at bivalent gene loci in melanoma. By preferentially binding promoters of genes marked by both activating and repressive histone modifications, TRIM24 appears to stabilize a poised chromatin state, limiting premature transcriptional activation of genes characteristic of the melanocytic states. Induction of TRIM24 expression alters the chromatin landscape of such bivalent genes which may serve as a trigger for rapid melanoma cell state transitions.

TRIM24 has been described as an oncogenic transcriptional activator of the androgen receptor (AR) in prostate cancer (Groner et al., 2016) while it acts as a co-repressor complex that suppresses murine hepatocellular carcinoma (Herquel et al., 2011), highlighting cell-type specific functions that may rely on differential interactions with specific TF and co-factors, either co-activators and/or repressors. DNA-binding motifs enrichment analysis on TRIM24-occupied sites identified a co-location with the well-known AP1 complex, suggesting a putative cooperative effect. While TRIM24 has been shown to interact with nuclear receptors (AR, ERα) (Tsai et al., 2010; Groner et al., 2016), STAT3 (Lv et al., 2017) or p53 (Allton et al., 2009; Isbel et al., 2023) in carcinoma models, our data suggest that it could potentially interact and cooperate with other transcription factors (such as AP1 or the Sp1/KLF family) in a melanoma-specific context. Furthermore, the motifs enriched in the vicinity of TRIM24 binding seem modified depending on whether the genes are down-or up-regulated by the latter, suggesting that the recruitment of different co-activators or co-repressors through its protein-protein interaction domain regulate TRIM24 biological impact on transcriptional programs.

Interestingly, H3K23 acetylation, which is specifically recognized by TRIM24, has recently been associated to NCSC genes regulation and resistance to treatment in melanoma (Lu et al. 2024). Indeed, in NCSC melanoma cells which display a high ALDH1A3 activity, pyruvate-derived acetaldehyde is a source for histone H3-acetylation at specific NCSC gene loci, inducing a cancer stem cell-like state. This further suggests that TRIM24-mediated transcriptional regulation of NCSC cell state may be metabolically controlled through histone acetylation levels regulation.

Overall, our findings identify TRIM24 as a key regulator of resistance to targeted and immunotherapy in melanoma, likely through its role in the maintenance of mesenchymal-like cell states, previously associated with direct inhibition of anti-tumor immune response (Benboubker et al. 2022; Plaschka et al. 2022). TRIM24 emerges as a robust actor of tumor aggressiveness, with expression restricted to the tumor compartment, underscoring its potential as a selective and druggable target. In this respect, we validate the efficacy to revert resistance to targeted therapies of a proteolysis-targeting chimera (PROTAC), dTRIM24, designed to induce ubiquitin-mediated degradation of TRIM24. We further provide the proof-of-concept that TRIM24 inhibition may resensitize tumors to immune checkpoint blockade, thus making of TRIM24 a promising target in combination with current therapies.

## Materials and Methods

### Human tumor samples immunofluorescence analyses

Melanoma tumor samples were obtained through the Biological Resource Center of the Lyon Sud Hospital (Hospices Civils de Lyon) and were used with the patient’s written informed consent. This study was approved by a regional review board (Comité Scientifique et Éthique des Hospices Civils de LYON CSE-HCL – IRB 00013204, study n° 22_5680). 52 primary cutaneous melanoma patient lesions were used for multi-immunofluorescence analyses (Supplementary Table 1). Patients were stratified in relapsing (RFS<2 years) and non-relapsing (RFS>2years) based on quarterly disease extension assessments performed in standard clinical care.

3-µm tissue sections were cut from formalin-fixed paraffin-embedded human melanoma specimens. The sections underwent immunofluorescence staining using the OPAL™ technology (Akoya Biosciences) on a Leica Bond RX. DAPI was used for nuclei detection. TRIM24 (HPA043495, RRID: AB_10962915, Sigma-Aldrich), SOX10 (sc-365692, RRID: AB_1084400, Santa Cruz). Sections were digitized with a Vectra Polaris scanner (Perkin Elmer, USA). Using the Inform software (Perkin Elmer), an autofluorescence treatment of images was carried out and tissue segmentation was performed to identify epidermis, stroma and tumor. Cell segmentation was then applied to analyze the expression of each marker in each cell. The matrix of phenotype containing the X– and Y-positions of each cell as well as the mean nuclear intensities of each fluorescence staining was then further analyzed using the R software. Tumors were spatially reconstructed using the R plot() function.

### Cell culture and treatments

The A375 human melanoma cell line was purchased from ATCC and cultured in DMEM complemented with 10% fetal bovine serum (FBS) (Cambrex) and 100 U/ml penicillin-streptomycin (Invitrogen). In order to authenticate the cell lines, the expected major genetic alterations were verified by NGS sequencing. The absence of Mycoplasma contamination was verified every 3 weeks with the MycoAlert detection kit (Lonza). Previously described patient-derived C-09.10 cells, established from *BRAF^V600^* metastatic melanoma, was grown in RPMI complemented with 10% FBS and 100 U/ml penicillin-streptomycin (Durand et al., 2024). Previously described BrafV600-mutated Br25F4 and NRasQ61-mutated NR6.1 mouse melanoma cell lines were cultured in RPMI 1640 Glutamax (61870044, Life Technologies) complemented with 10% FBS (Cambrex) and 100 U/mL penicillin-streptomycin (15140148, Gibco) (Plaschka et al., 2022). The BRAF inhibitor PLX4032/vemurafenib was purchased from Selleck Chemicals (Houston, TX, USA) and reconstituted in DMSO. The PROTAC dTRIM24 (MedChemExpress) was reconstituted in DMSO at a stock concentration of 10 mM. Cells were treated for 14 days, with compound renewal every 3 days.

### Viral infections

For shRNA-Trim24 knock-down in human (A375 and C-09.10 cells) and murine (Br25F4 and NR6.1) melanoma cell lines, human embryonic kidney 293T cells (4 x 10^6^) were transfected with lentiviral expression constructs (10 µg) in combination with GAG-POL (5 µg) and ENV expression vectors (10 µg). The constructs allowed the insertion of shRNA targeting TRIM24 in a lentiviral plasmid vector pLKO.1-puro (MISSION, Sigma) for human (SHCLNG-NM_003852) and mouse (SHCLNG-NM_145076) or a non-target shRNA control plasmid (SHC016-1EA). Viral supernatants were collected 48 h post-transfection, filtered (0.45 µm membrane), and placed in contact with 2 x 10^6^ melanoma cells for 8 h in the presence of 8 µg/mL polybrene. The cells were selected in the presence of puromycin (1 µg/mL) (Invitrogen) 48h post infection.

### Immunoblot analyses

Cells were washed twice with phosphate buffered saline (PBS) containing CaCl_2_ and then lysed in a 100 mM NaCl, 1% NP40, 0.1% SDS, 50 mM Tris pH 8.0 RIPA buffer supplemented with a complete protease inhibitor cocktail (Roche, Mannheim, Germany) and phosphatase inhibitors (Sigma-Aldrich). A Trim24 antibody was used (Sigma-Aldrich, HPA04349). Loading was controlled using anti-GAPDH (Sigma-Aldrich, Cat# ABS16 (RRID: AB10806772). Horseradish peroxidase-conjugated goat anti-rabbit polyclonal antibodies (Glostrup) were used as secondary antibodies. Western blot detections were conducted using the Luminol reagent (Santa Cruz). Western Blot Digital Imaging was performed with the ChemiDoc™ MP Imager (Bio-Rad).

### RT-Q-PCR

Total RNA was isolated using the RNeasy Kit (QIAGEN) and reverse-transcribed using a high cDNA capacity reverse transcription kit (Maxima First Strand cDNA synthesis Kit, Thermoscientific) following the manufacturer’s instructions using 1000 ng of RNA as a reverse transcription template in a 20 μL final volume. The samples were incubated for 10 minutes at 25°C, followed by 15 minutes at 50°C and 5 minutes at 85°C in T100 Thermal Cycler (1861096, Bio-Rad). Real-time qPCR reactions were performed using One*Green*® FAST qPCR Premix (OZYA008-200XL, Ozyme) according to the manufacturers protocol. Reactions were done using 15ng of cDNA template and 1µM of each primer. All reactions, including no-template controls and RT controls were performed in triplicate on an Azure Ciel real Time PCR (Ozyme) pre-denaturation 3 minutes at 95°C then 40 cycles at 95°C for 5s followed by 30s at 60°C. Results were analyzed with the Azure Cielo manager software. Human GAPDH was used for normalization. Exon-spanning probes were designed using the primer-blast (NCBI).

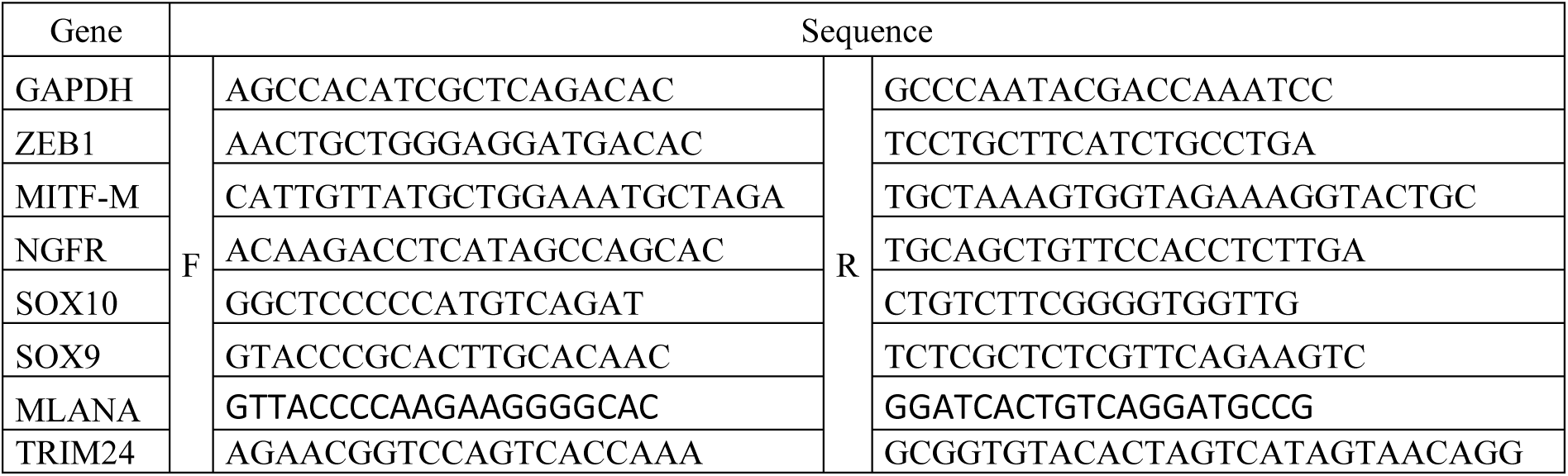

### Wound healing assays

Wound healing assays were performed using incucyte assay (Sartorius). Briefly, A375 cells were trypsinized and seeded at 60000 cells per well in a 96 well-plate. The day after, the cells were treated with 10 µg/ml of mitomycin and 2h after the scratch module was used to create the wound. EssenBioScience IncuCyte Zoom System was used to acquire images and measure the wound in real-time every 2 h for 48h.

### Rezasurin and Incucyte viability assays

Viability was measured using rezasurin assays. 500 A375 and 2000 C-09.10 cells were plated per well in 96-well plates. Cells were treated for 7 days and the medium was renewed 3 times a week with indicated PLX4032 concentration (Selleckchem) and the same DMSO volume for control. The relative viability was assessed by measuring resorufin fluorescence on the ClarioSTAR plate reader (BMG LABTECH).

For incucyte assays, 2 000 A375 and 20 000 C-09.10 cells were seeded onto a 24-well plate. After 24 h, cell medium was renewed with indicated treatments, as well as PLX4032 and propidium iodide (Sigma-Aldrich, 1/3000). The EssenBioScience IncuCyte Zoom Live-Cell Analysis System was used to measure and analyze real-time cell mortality every 2 h. Dead cells were marked with propidium iodide. Data were then converted into prism in order to draw graphs.

### Sphere formation assay

Sphere formation assay was performed by seeding 5, 10 and 20 A375 cells per well of a 96 low adherence culture plate in DMEM-F12 medium supplemented with FGF7 (5 ng/mL, Peprotech), FGF10 (20 ng/mL, Peprotech), FGF10 (20 ng/mL, Peprotech), and 1X B27 (Gibco). The medium was renewed two times a week. The sphere were allowed to grow for 3 weeks and images were acquired with EssenBioScience IncuCyte Zoom Live-Cell Analysis System to determine the number of well containing a melanoma sphere.

### RNA sequencing

RNA libraries were prepared with the TrueSeq poly-A+ kit from Illumina and sequenced on the genomic platform of the CRCL, on an Illumina NovaSeq 6000 sequencing machine with a paired-end protocol (2×75bp, 32Mp reads).

### CUT&Tag sequencing

CUT&Tag assays were carried out using iDeal CUT&Tag Kit for Histones for chromatin profiling (Diagenode) following manufacturer’s recommendations. Each condition was performed on 150 000 cells. Chromatin fragments were immunoprecipitated with antibodies directed against TRIM24 (1 μg, BETHYL-A300-815) or IgG (1 µg, Bio-Rad, PRABP01, RRID: AB321631) as negative control. Antibodies targeting H3K4me3 (C15200152), H3K27me3 (C15200181-50), H3K23Ac (C15410344), H3K27Ac (C15200184-50), H3K4un (C15200149), H3K9Me (C15410193) were all CUT&Tag validated and acquired from Diagenode. Immunoprecipitated DNA was purified and dissolved into 25 µl of DNA elution buffer. DNA libraries were prepared with the Diagenode MicroPlex Library Preparation Kit v2, and sequenced on an Illumina Novaseq sequencing machine (paired-end protocol, 75bp, 80 M reads) on the genomic platform of the CRCL. The library was verified using Bioanalyzer (Agilent). Two biological replicates from two independent experiments were performed for each CUT&Tag condition.

### ATAC sequencing

ATAC-seq was performed using ATAC-seq kit for open chromatin assessment kit (Diagenode) following manufacturer’s instructions. Each condition was performed in duplicate using 1.5 x 10^5^ cells as an input. DNA libraries were prepared with the Diagenode MicroPlex Library Preparation Kit v2, and sequenced on an Illumina Novaseq sequencing machine (paired-end protocol, 75bp, 80 M reads) on the genomic platform of the CRCL. The library was verified using Bioanalyzer (Agilent). Two biological replicates from two independent experiments were performed for each ATACSeq condition.

### Mouse injections

Experiments using mice were performed in accordance with the animal care guidelines of the European Union and French laws and were validated by the local Animal Ethic Evaluation Committee and the French MESRI (APAFIS #43229; APAFIS #46228; APAFIS #38245). Mice were housed and bred in a specific pathogen-free animal facility “AniCan” at the CRCL, Lyon, France. Single cell suspensions of Br25F4 and NR6.1 cell models (1-3 x 10^6^ cells), in PBS/Matrigel (BD Biosciences, Oxford, UK) (1:1) were injected subcutaneously into the flank of six-week-old male C57BL/6J mice (Charles River laboratories). 1-3 x 10^6^ A375 cells were injected subcutaneously into the flank of six-week-old female Swiss Nude (Charles River laboratories). n=5 mice were included in each experimental group, in separate cages. No randomization was performed. Tumor growth was monitored for 2-6 weeks post-injection. Tumors grew up to 1.5 cm in diameter, at which point animals were euthanized. For anti-PD-1 treatment, 5 days after injection, mice were treated with intra-peritoneal injection of 200 µg of anti-PD-1 rat anti-mouse PD-1 clone RMP1-14 (BP0146, Bio X Cell) or with the control isotype 3 times every 2 days.

### IHC staining analyses of mouse tumor samples

Tumors were embedded in paraffin and TRIM24 staining was performed using the anti-TRIM24 antibody (Sigma-Aldrich, HPA04349) and purple chromogen (for heavily pigmented tumors) detection and counterstaining with hematoxylin. Images were digitalized with a 3DHistech Pannoramic SCAN2 scanner on the Research Pathology Platform. Quantification was done with HALO™ Image Analysis Software (Indica Labs).

## Data availability

The data reported in this paper have been deposited in the Gene Expression Omnibus (GEO) database under accession number GSE297251.

Public data were from: Bulk RNA-seq data of melanoma cell lines from (Tsoi et al., 2018) were retrieved from GEO, with accession number GSE80829. Single-cell RNAseq data of melanoma cell lines from (Wouters et al., 2020) were retrieved from GEO (GSE134432) and from (Pozniak et al., 2024) and (Karras et al., 2022) from the KU Leuven Research Data Repository. TCGA data of melanoma tumors in cluding RSEM-normalized RNseq data were analysed and retrieved from cbioportal.

## Statistical analyses and data visualization

To ensure adequate power and decreased estimation error, we performed multiple independent repeats and experiments were conducted at least in triplicate. Data are presented as mean ± s.d. or ± s.e.m as specified in the figure legends. Statistical analyses were performed using GraphPad Prism 10 software (GraphPad Software, Inc., San Diego, USA) or R software (v4.3.2) (R Core Team, 2021. All statistical tests were two-tailed and p-values were corrected, when indicated, with the Benjamini-Hochberg method. Unpaired Student’s t-tests or Mann– Whitney tests were used to compare the means of two groups, as specified in the figure legends. Graphs were produced with either base R functions or the ggplot2 package (v3.5.1). All heatmaps were generated using the R package ComplexHeatmap (v2.18.0) with Ward.D clustering method and Euclidean distance. Venn diagrams were conducted with *eulerr* (v7.0.1) and VennDiagram (v1.7.3) R package.

## Author contributions

SDu and FB designed, performed, analyzed experiments and prepared figures. SDu cosupervised the project with JC and wrote the manuscript. MG and LB performed and analyzed experiments. FP and MD contributed to multi-IF analyses. RP, EC, RS and AF performed RNA-Seq, Cut & tag and ATAC-seq bioinformatics analyses. SDa and AE provided human samples, clinical data and funding and wrote the manuscript. JC conceived and supervised the whole project and wrote the manuscript.

## Acknowledgments

The authors would like to thank Brigitte Manship for critical reading, Zdenko Herceg team for useful discussions, Thibault Andrieu for cytometry analyses. This work was funded by the Ligue Nationale contre le Cancer (Comité de l’Ain), the Lyon Integrated Research Institute in Cancer (SIRIC LYriCAN INCa-DGOS-Inserm_12563 and LYriCAN+ INCa-DGOS-INSERM-ITMO cancer_18003), the Institut Convergence PLAsCAN (ANR-17-CONV-0002), the ERiCAN program of Fondation MSD-Avenir (Reference DS-2018-0015), the Institut National contre le Cancer (INCA-DGOS PRTK), ARC sign’it 2019, the Société Française de Dermatologie (SFD), the association Melarnaud and Vaincre le Mélanome. FB was supported by a fellowship from ITMO Cancer of Aviesan within the framework of the 2021-2030 Cancer Control Strategy, on funds administered by Inserm. SD was supported by a fellowship from the Association pour la Recherche contre le Cancer (ARC).

## Notes

### Competing Interest Statement

The authors have declared no competing interest.

