## Supplementary Figures and Methods for "The epigenetic regulator TRIM24 controls melanoma cell dedifferentiation and resistance to treatment in melanoma"

### Supplementary Figure 1

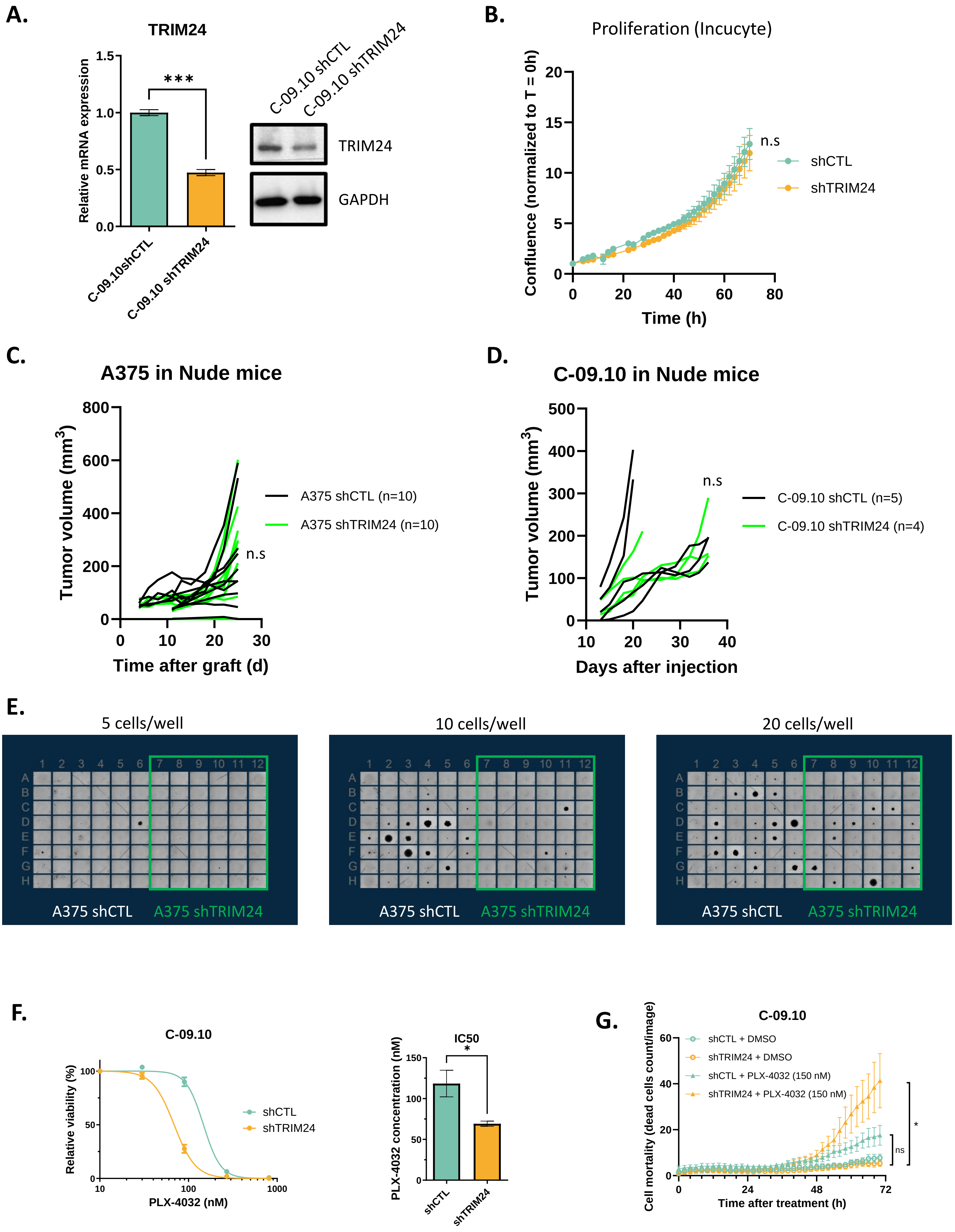

**Supplementary Figure 1: *TRIM24* knock-down does not impair cell proliferation nor tumor growth but impacts sphere formation ability.**

**A.** Evaluation of shRNA knockdown of *TRIM24* by RT-qPCR (left) and Western-blot (right) in C-09.10 melanoma cells. **B.** Proliferation of A375 cell line upon *TRIM24*-knock-down (KD) measured by incucyte assay. **C-D.** Tumor growth of A375-sh*TRIM24* **(C)** and C-09.10-sh*TRIM24* **(D)** cell lines after xenograft in Nude mice. **E.** Microscopy images of sphere formation in A375 cells upon *TRIM24*-KD. **F.** Viability curve (left) and IC50 values (right) for BRAFi PLX4032 in C-09.10 cells after 6 days of treatment upon *TRIM24*-KD. **G.** Measure of C-09.10 cells mortality over-time measured incucyte assay comparing sh-*TRIM24* and shCTL cells, with and without PLX4032 treatment at 150nM.

### Supplementary Figure 2

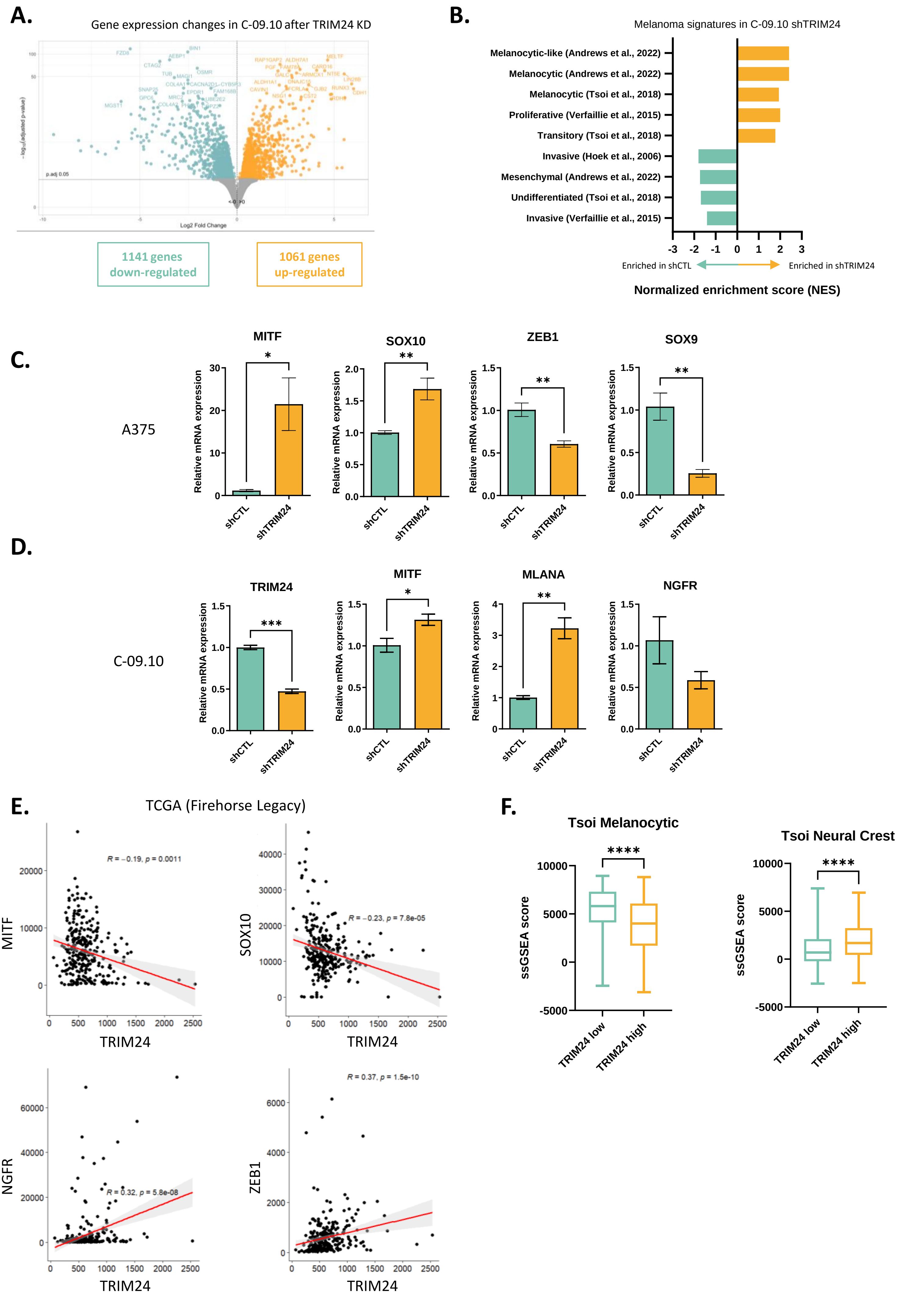

**Supplementary Figure 2: TRIM24 mediates melanoma cell dedifferentiation towards a NCL/invasive phenotype**

**A.** Volcano plot of differentially expressed (DE) genes by RNA-seq in C-09.10 cells after TRIM24-knock-down (KD). The number of genes up- and down-regulated are indicated. **B.** Normalized enriched scores (NES) upon TRIM24-KD in C-09.10 of various already described melanoma cell phenotypes signatures. **C-D.** Relative expression measure by RT-qPCR of several well-defined melanoma cell markers upon TRIM24-KD in A375 (**C**) and C-09.10 (**D**). **E.** Correlation of TRIM24 expression with MITF, SOX10, NGFR and ZEB1 in the TCGA database. **F.** Expression of the melanocytic and neural-crest-like signatures from Tsoi et al in TRIM24low and TRIM24high tumors in the TCGA cohort.

Supplementary Figure 3

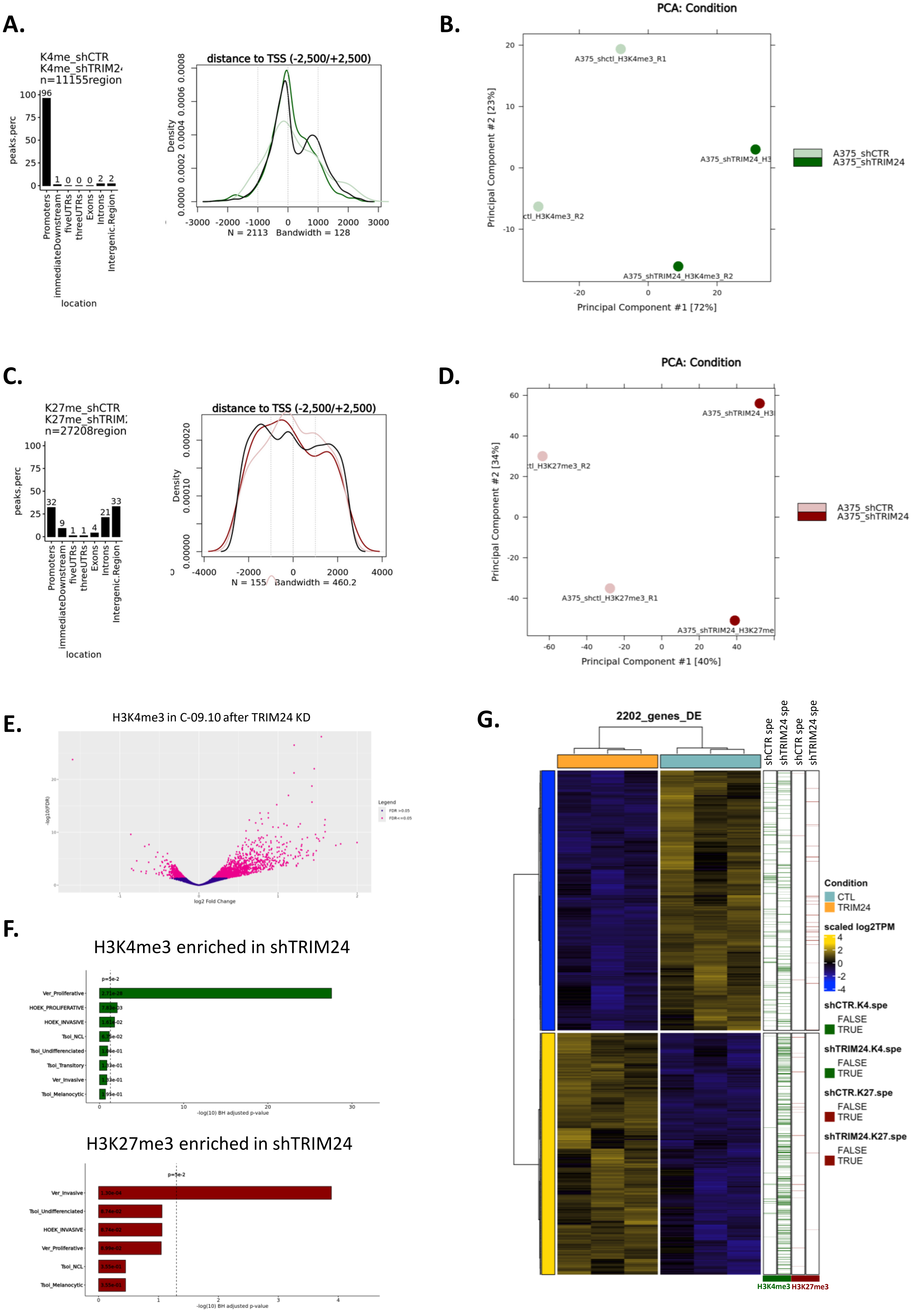

**Supplementary Figure 3: TRIM24 mediates activating and repressive histone marks remodelling associated with alterations in transcriptional programs**

**A.** Localization of H3K4me3 marks and their distance to TSS (right) in A375 cells . **B.** Principal component analysis (PCA) on H3K4me3 histone marks in A375 shCTL and sh*TRIM24*. Each replicate is plotted. **C.** Localization of H3K23me3 marks in A375 cells. **D.** Principal component analysis (PCA) on H3K27me3 histone marks in A375 shCTL and sh*TRIM24*. **E.** Volcano plot displaying the differential binding of the activating H3K4me3 histone mark associated to a gene in C-09.10 sh*TRIM24* versus shCTL. **F.** Melanoma cell signatures enrichment in genes presenting a gain of these histone marks in sh*TRIM24* versus shCTL, with H3K4me3 (top) and H3K27 (bottom). **G.** Heatmap of DEs genes upon sh*TRIM24* in C-09.10 cells. H3K4me3 and H3K27me3 histone marks specific to sh*TRIM24* or shCTL are indicated.

### Supplementary Figure 4

**A.**

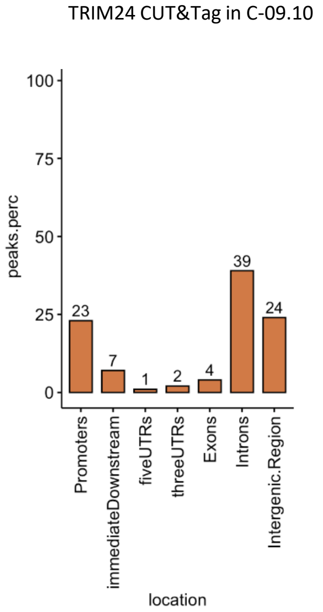


**B.**

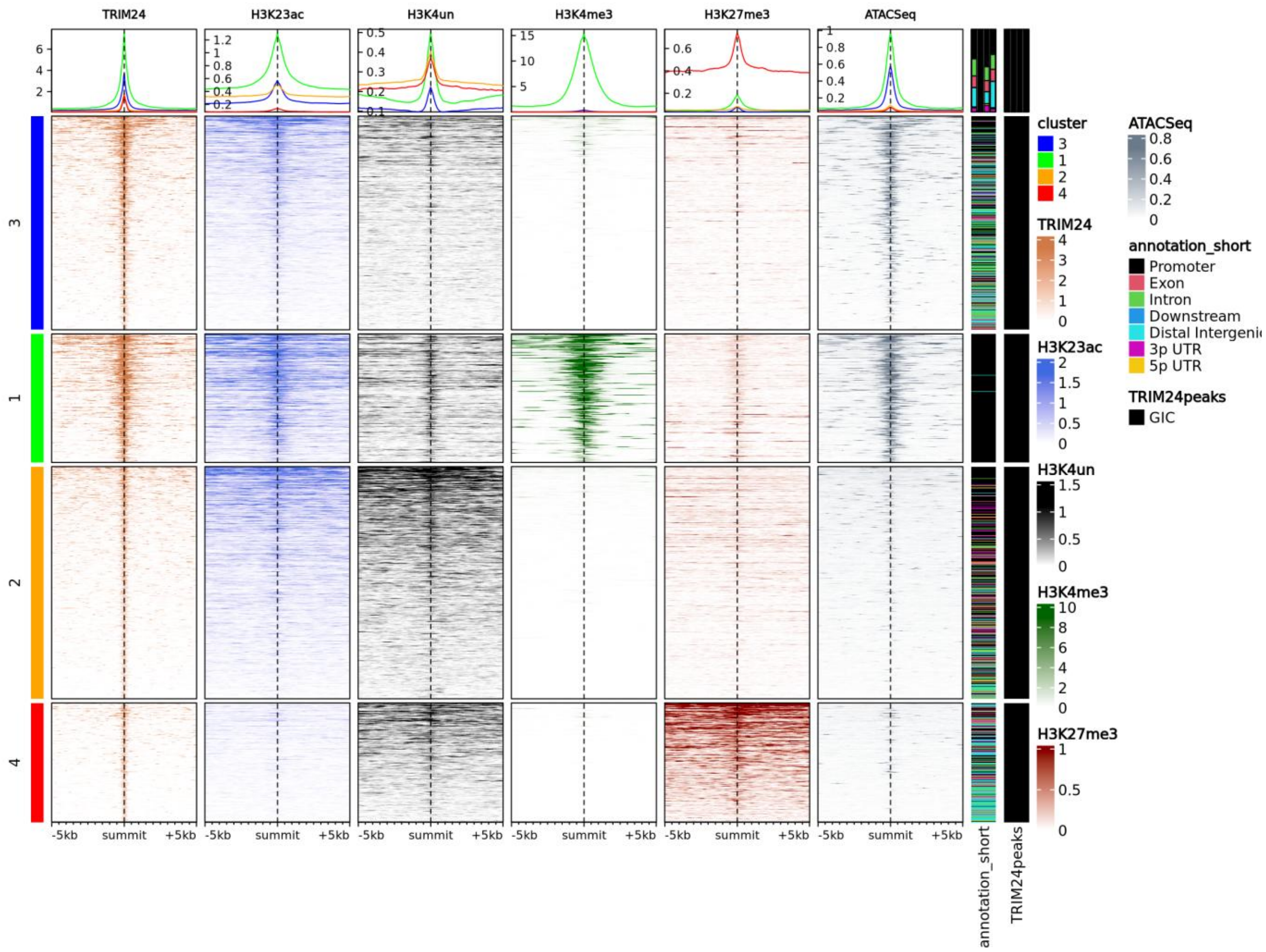


**C.**

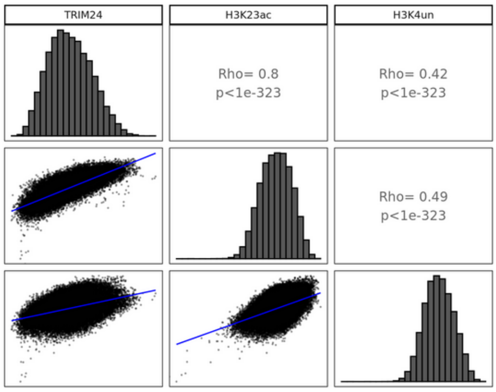

**Supplementary Figure 4: TRIM24 binds H3K23ac marks and open chromatin in genes related to melanoma cell phenotype**

**A.** Localisation of TRIM24 peaks in the genome in C-09.10 cells identified by TRIM24 Cut&Tag analysis. **B.** Read density heatmap of the peak profiles for histone marks (H3K23ac, H3K4un, H3K4me3 and H3K27me3) relative to summits of TRIM24 peaks and chromatin accessibility (ATAC-seq) in C-09.10 cell line. The profiles are clustered using hierarchical clustering method, the profile intensity of each mark in each cluster is indicated. Genome annotations are indicated. **C.** Correlogram studying the association between TRIM24 and H3K23ac and H3K4un peaks.

### Supplementary Figure 5

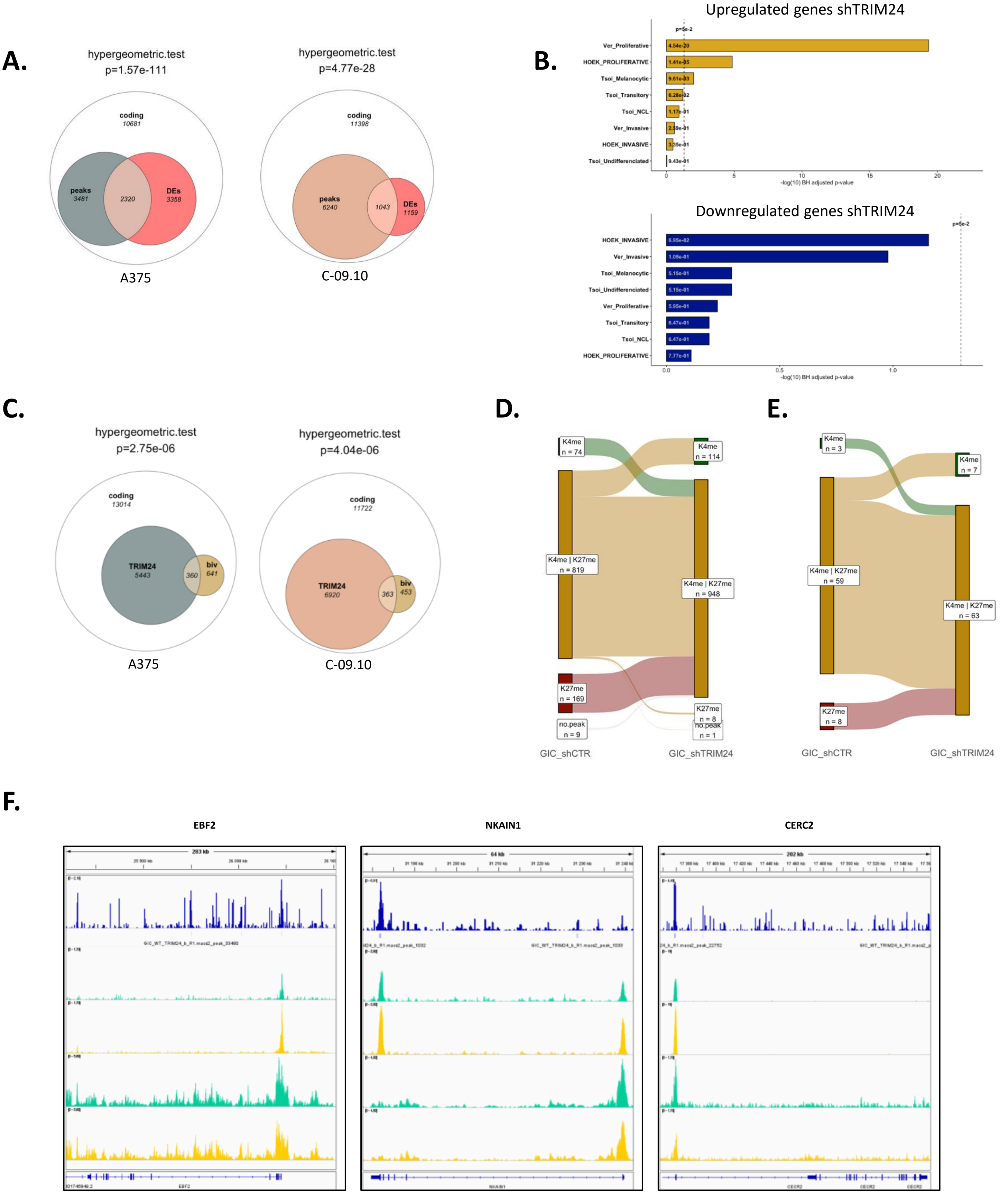


G.

| Rank | Motif | P-Value | % of targets | % of background | Corresponding TF |
|---|---|---|---|---|---|
| 1 | 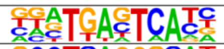 | 1e-125 | 9.69% | 3.52% | AP-1 |
| 2 | 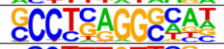 | 1e-70 | 26.90% | 18.43% | Tfap2C |
| 3 | 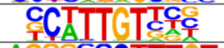 | 1e-68 | 18.49% | 11.48% | Sox10 |
| 4 | 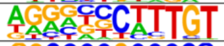 | 1e-56 | 10.96% | 6.08% | Sox9 |
| 5 | 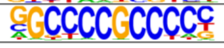 | 1e-53 | 12.60% | 7.43% | Sp1 |

**Supplementary Figure 5: TRIM24 binding at bivalent genes modifies chromatin state**

**A.** Venn diagram displaying the proportion of TRIM24 binding at differentially expressed genes upon *TRIM24*-KD in A375 and C-09.10 cell lines. The statistically significant enrichment was assessed by an hypergeometric test. **B.** Enrichment of melanoma cell phenotype signatures on genes associated to a TRIM24 peak and differentially expressed upon *TRIM24*-KD in C-09.10 cells. **C.** Venn diagram displaying the proportion of TRIM24 binding at bivalent gene promoters in A375 and C-09.10 cell lines. The statistically significant enrichment was assessed by an hypergeometric test. **D-E**. Effect of *TRIM24* loss on genes presenting a bivalent status in C-09.10 cell lines. Evolution of H3K4me3 and H3K27me3 histone marks patterns upon *TRIM24*-KD on all bivalent genes (**D**) and bivalent genes composing the melanoma proliferative signature (**E**) (Verfaillie et al., 2015). **F.** IGV visualisation of EBF2, NKAIN1 and CERC2 loci presenting a bivalent status. The binding of TRIM24 as well as H3K4me3 and H3K27me3 signals upon *TRIM24*-KD are included. Two replicates are overlayed for each condition. **G.** Top 5 HOMER-identified enriched motifs at TRIM24 binding sites in C-09.10 cells. The associated p-values and the percentages of motif representation on target and background signal are indicated.

### Supplementary Figure 6

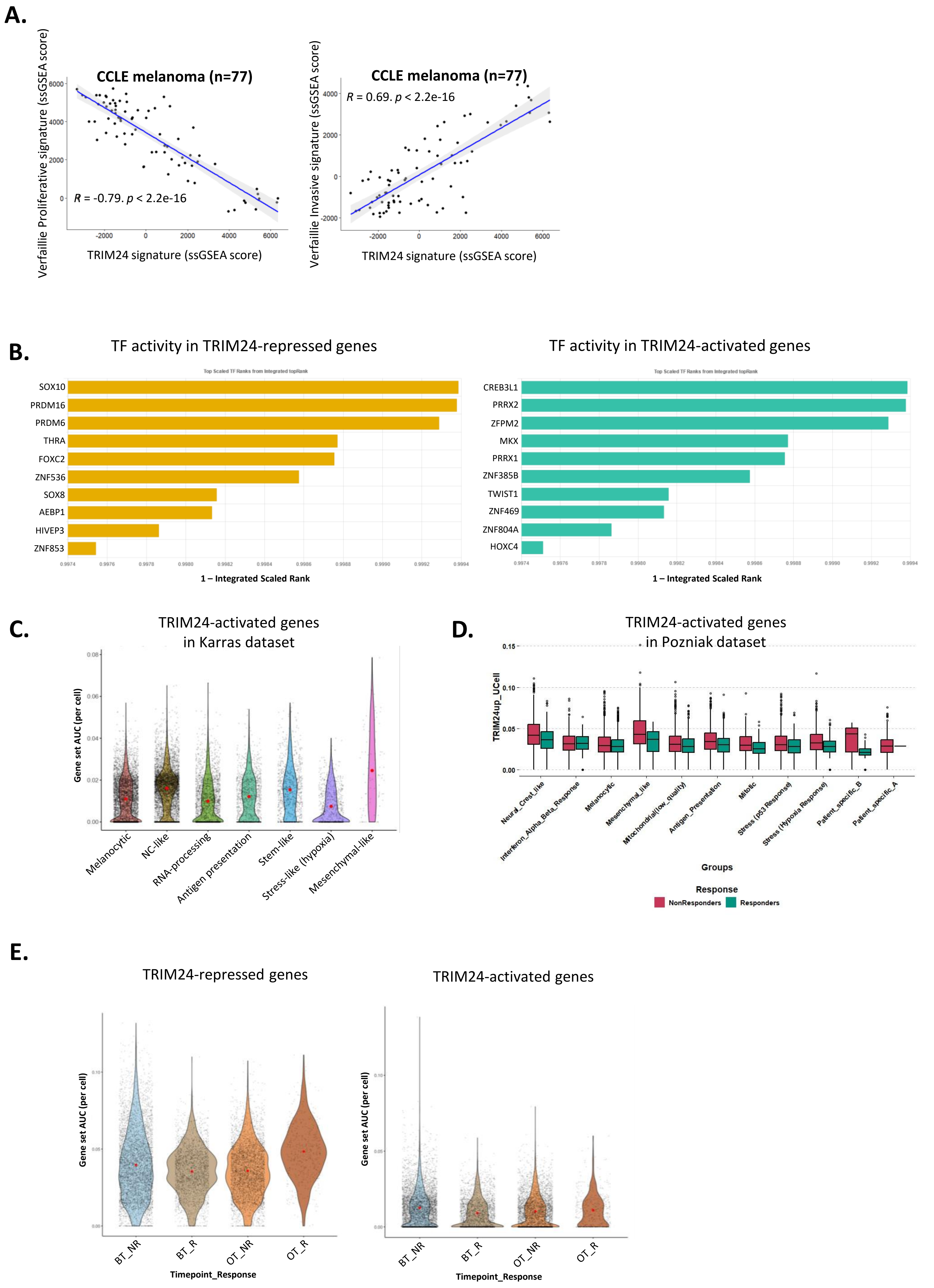

**Supplementary Figure 6: TRIM24 signature is enriched in mesenchymal and NCL therapy-resistant melanoma cell states.**

**A.** Correlation of TRIM24 signature with proliferative and invasive signatures in CCLE melanoma RNAseq dataset from Verfaillie *et al.* **B.** TF activity enriched in genes downregulated (left) and upregulated (right) genes from TRIM24 signature. **C.** Violin plot on sc-RNAseq data showing the TRIM24-activated genes expression in the from Karras *et al*. dataset. **D.** TRIM24 signature in the distinct melanoma subpopulations from Pozniak *et al*. according to patients ICB response status. **E.** Up- and down-regulated genes from TRIM24 signature in ICB responders and non-responders before (BT) and on (OT) immunotherapy treatmentfrom Pozniak *et al.*

### Supplementary Figure 7

**A.**

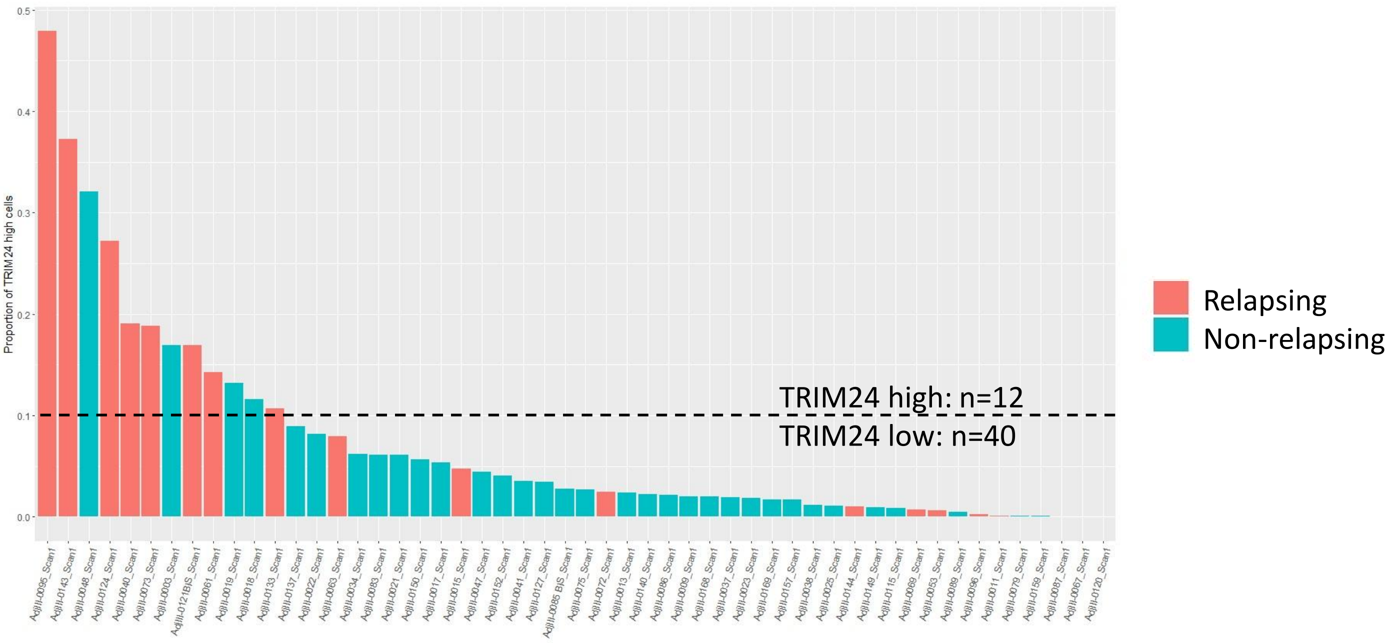


**B.**

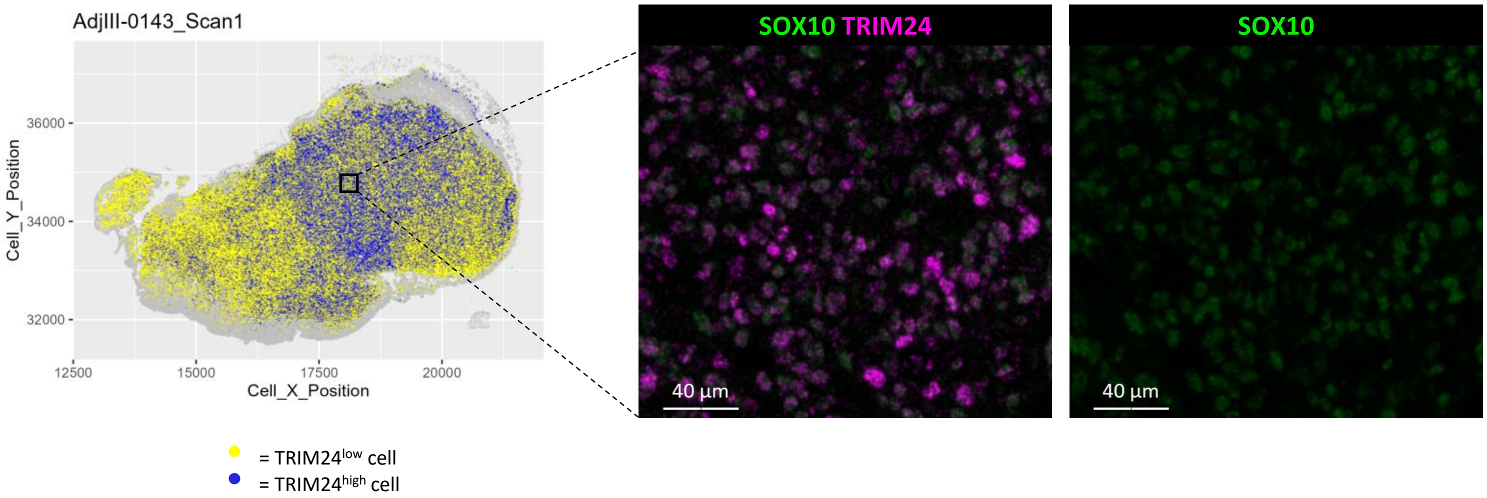


**C.**

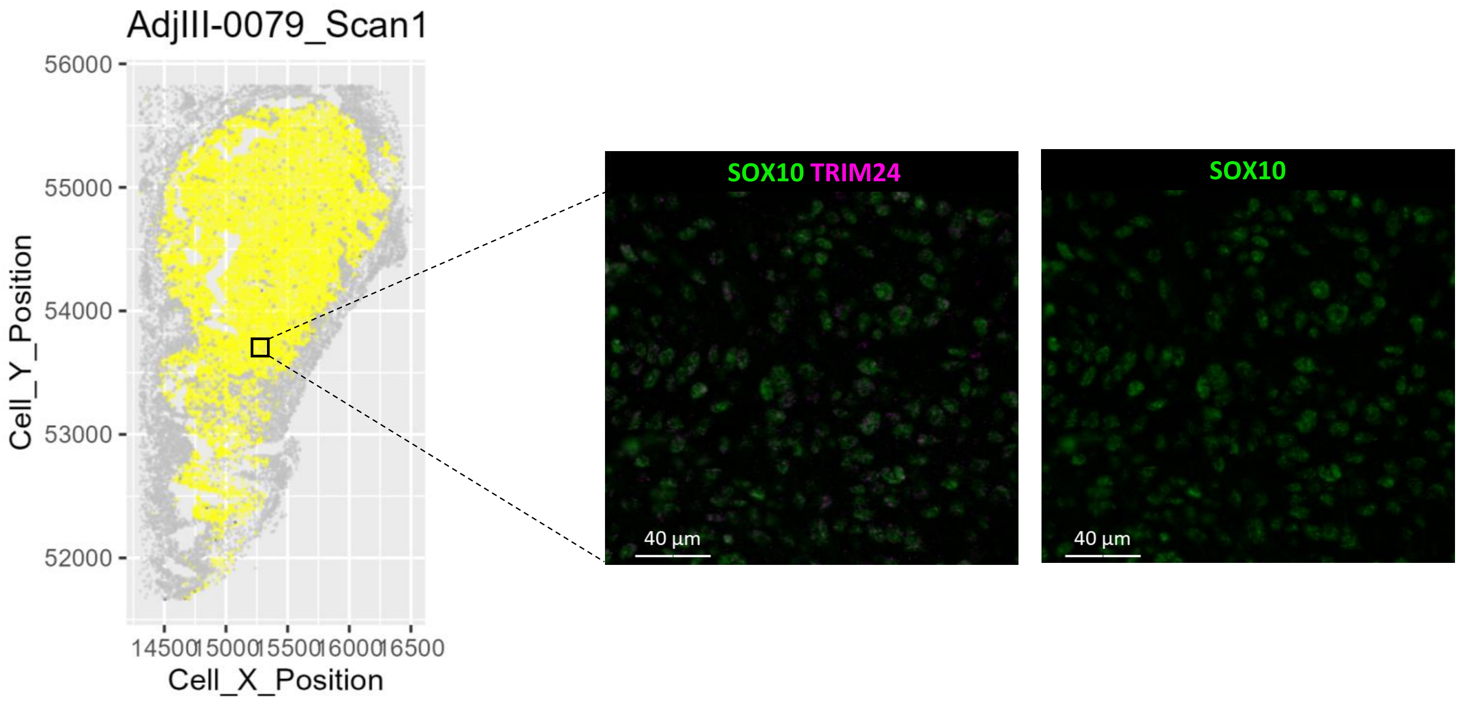

**Supplementary Figure 7: TRIM24 expression heterogeneity in patients melanoma tumors**

**A.** Barplot representing the proportion of melanoma cells with a high expression of TRIM24 in the melanoma cohort. A cut-off of 10% of TRIM24$^{high}$ was defined to segregate TRIM24$^{high}$ versus TRIM24$^{low}$ tumors. The response status to immune checkpoint blockade is indicated: relapsing (pink), non-relapsing (blue). **B-C**. Spatial reconstitution (left) and representative multiplex-IF images (right) of a TRIM24high **(B)** and a TRIM24 low **(C)** tumor.

### Supplementary Table 1

**Clinical parameters of stage III melanoma patients treated with anti-PD1.**

| | **Non-relapsing**<br>**N = 35 (67.3%)** | **Relapsing**<br>**N = 17 (32.7%)** | ***Statistic*** |
|---|---|---|---|
| **Age** | | | t-test, ns |
| Years, mean +/- SD | 63.54 +/- 15.0 | 67.1 +/- 16.2 | |
| **Sex** | | | Ficher's exact test, ns |
| Female, n (%) | 11 (31.4%) | 7 (41.2%) | |
| Male, n (%) | 24 (68.6%) | 10 (58.8%) | |
| **BRAF/NRAS mutational status** | | | Ficher's exact test, ns |
| BRAF, n (%) | 13 (37.1%) | 8 (47.1%) | |
| NRAS, n (%) | 12 (34.3%) | 4 (23.5%) | |
| Non-mutated, n(%) | 8 (22.9%) | 5 (29.4%) | |
| ND, n(%) | 3 (8.6%) | 0 | |
| **Breslow index** | | | Mann-Whitney, p = 0.0453 |
| mm, median (range) | 4.00 (1.0-18.0) | 6.50 (1.2-19.0) | |
| **Ulceration** | | | Ficher's exact test, ns |
| Yes, n (%) | 19 (54.3%) | 12 (70.6%) | |
| No, n (%) | 16 (45.7%) | 4 (23.5%) | |
| ND, n (%) | 0 | 1 (5.9%) | |

ND: not determined; SD: standard deviation; Relapse status 1 year after treatment.

**Supplementary methods**

**RNA sequencing analyses**

Sequencing control metrics were computed using FastQC (v0.11.9) (LaMar, 2015). For careful QC and batch effect analyses, raw data were aligned on the human genome (GRCh38) with STAR (v2.7.8a), and RNA control metrics were evaluated using RSeQC (v4.0.0) (Dobin 2013 ; Wang 2012). Gene expression was quantified with Salmon (1.4.0) on the raw sequencing reads, using gencode v37 comprehensive annotation set (Patro 2017). Further gene expression analyses were restricted to protein-coding genes using *annotables* R package (v.0.2.0) (protein coding, immunoglobulins (IG) and T cell receptors (TR) biotypes) (Turner 2024).

Differential expression analyses were performed using the R package *DESeq2* (v1.42.0) (Love 2014), with Wald test and apeglm shrinkage estimator (v1.24.0) **(**Zhu 2019**)**.

**scRNA Sequencing analysis**

Single-cell RNAseq data were retrieved as a Seurat Object, the data were analysed and visualized using Seurat (4.3.0) and SCpubr (1.1.2) packages. The data normalization, data filtering, dimensional reductions and cell annotations are the same as used by the authors (Wouters and Pozniak). Signatures scores were calculated using the UCell R package (v1.2.1).

**CUT&Tag sequencing analysis**

The nfcore/ cutandrun analysis pipeline (v3.2.1) was used to process the data (https://github.com/nf-core/cutandrun) (Meers 2019). BigWig coverage files were generated, and peak calling was performed from BAM files using MACS2 (v2.2.7.1) (Zhang et al., 2008) for narrow peaks such (e.g., TRIM24, H3K4me3 and H3K27ac) or using EPIC2 (v0.0.52) (Stovner and Sætrom, 2019) for broad marks (e.g., H3K4un, H3K9me3, H3K23ac and

H3K27me3). Peak calling was conducted either independently for each sample (for differential binding and motif enrichment analyses), or on merged replicates (for bivalent status assignment, signal correlation and density heatmap visualization), using the corresponding IgG immunoprecipitation as a control.

**ATAC sequencing analysis**

The nfcore/atacseq analysis pipeline was used to process the data (https://github.com/nf-core/atacseq). Briefly, after adapter trimming using Trimgalore, fastq files were aligned with BWA to the human reference genome GRCh38. Reads were then filtered out in order to avoid blacklisted regions (form ENCODE), duplicates, unmapped, multiple locations, >4 mismatches, insert size >2kb, different chromosomes and other than FR orientation mappings. Normalized BigWig (scaled to 1 million mapped reads) were generated and peak calling was performed with MACS2.

**Peaks annotation and analysis**

For both ATAC-seq and CUT&Tag-seq, genomic localization of called peaks was performed through *assignChromosomeRegion* function from *ChIPpeakAnno* R package (v3.36.1). Gene annotation data were obtained from *TxDb.Hsapiens.UCSC.hg38.knownGene* R package (v3.18.0) using "transcript" as feature. Distance to closest TSS was defined with *annotatePeakInBatch* function with the output set as "nearestLocation”. Peak-to-gene assignment was conducted through the *annotatePeakInBatch* function using the following options: output="overlapping", FeatureLocForDistance="TSS", select="all". For narrow peaks (e.g., TRIM24, H3K4me3, and H3K27ac), binding regions were defined as -1000 to +500 bp relative to the TSS, whereas for broad peaks (e.g., H3K4un, H3K9me3, H3K23ac, and H3K27me3), regions extended from -2500 to +500 bp. A bivalent status was assigned to each gene showing both an H3K4me3 peak within the -1000 to +500 bp region relative to the TSS, and an H3K27me3 peak within the -2500 to +500 bp region. Differential analyses were performed using *Diffbind* (v3.12.0) and DESeq2 (v1.42.0) R packages (Starck 2011, Love 2014). Motif enrichment analysis was conducted employing *findMotifsGenome* function, from HOMER software (v4.11.1) (Heinz et al., 2010).

The motifs were searched 400-bp regions centered on each peak summit. Read density heatmaps, signal correlation (rowmeans) and clustering analysis were performed with *profileplyr* R package (v1.18.0).

**Pathway enrichment and score analysis**

Previously published melanoma signatures were retrieved from Tsoi et al 2018, Andrews et al 2022, Pozniak et al 2024, Belote et al 2021, Verfaillie et al 2015.

For both RNA-seq and CUT&Tag-seq analyses, overrepresentation of lists of genes in specific biological pathways were tested using *clusterProfiler* (v4.10) R package (Yu 2012).

For RNA-seq, Gene Set Enrichment Analyses (GSEA) were carried out using fgsea R package (v1.28.0) and gene lists were pre-ranked using Signal2Noise metric.

Single sample GSEA (ssGSEA) scores were computed on TPM normalized data with *gsva* R package (v10.1.x) (Hänzelmann 2013).
